# A wide diversity of viruses detected in African mammals involved in the wild meat supply chain

**DOI:** 10.1101/2024.11.28.625648

**Authors:** Mare Geraerts, Sophie Gombeer, Casimir Nebesse, Douglas Akaibe, Dudu Akaibe, Pascal Baelo, Anne-Lise Chaber, Guy-Crispin Gembu, Philippe Gaubert, Léa Joffrin, Anne Laudisoit, Nicolas Laurent, Herwig Leirs, Claude Mande, Joachim Mariën, Steve Ngoy, Jana Těšíková, Ann Vanderheyden, Rianne van Vredendaal, Erik Verheyen, Sophie Gryseels

**Affiliations:** Evolutionary Ecology group, Department Biology, University of Antwerp, Wilrijk, Belgium; Swiss Federal Institute for Forest, Snow and Landscape Research WSL, Birmensdorf, Switzerland; OD Taxonomy and Phylogeny, Royal Belgian Institute of Natural Sciences, Brussels, Belgium; University of Kisangani, Kisangani, Democratic Republic of the Congo; School of Animal and Veterinary Sciences, University of Adelaide, Adelaide SA, Australia; Centre de Recherche sur la Biodiversité et l’Environnement (CRBE), Université de Toulouse, CNRS, IRD, Toulouse INP, Université Toulouse 3 – Paul Sabatier (UT3), Toulouse, France; CIIMAR/CIMAR, Interdisciplinary Centre of Marine and Environmental Research, University of Porto, Terminal de Cruzeiros Do Porto de Leixões, Avenida General Norton de Matos s/n, 4450-208 Matosinhos, Portugal; Ecohealth Alliance, 520 8th Avenue Room 1200, 10018 New York, USA; OD Taxonomy and Phylogeny (BopCo), Royal Belgian Institute of Natural Sciences, Brussels, Belgium; Department of Environmental Sciences, Open Universiteit, Heerlen, The Netherlands; Virus Ecology group, Department of Biomedical Science, Institute of Tropical Medicine Antwerp, Antwerp, Belgium

## Abstract

The processes involved in acquiring, trading, preparing, and consuming wild meat pose significant risks for the emergence of zoonotic infectious diseases. Several major viral outbreaks have been directly linked to the wild meat supply chain, yet our knowledge of the virome in many mammals involved in this chain remains limited and disproportionately focused on certain mammalian taxa and pathogens.

This report presents the findings of a metagenomic viral screening of 99 specimens belonging to 27 wild African mammal species and one domesticated species, all traded for their meat. The study focuses on tissue and swab samples collected from various regions in the Democratic Republic of the Congo and in Brussels, Belgium.

A total of fifteen virus strains were detected, belonging to the families *Arteriviridae*, *Retroviridae* and *Sedoreoviridae* (primates), *Picobirnaviridae* (primates and rodents), *Picornaviridae* (rodents), *Hepadnaviridae* (hyrax), *Orthoherpesviridae* (artiodactylid and carnivore) and *Spinareoviridae* (carnivore). Several strains were detected in mammalian hosts for the first time, expanding their host range and genetic diversity. Of note is the presence of viruses genetically related to recognised zoonotic pathogens, i.e., human picobirnavirus (*Orthopicobirnavirus hominis*) (primates and rodents), simian foamy viruses (*Simiispumavirus*) (primates), and rotavirus A (*Rotavirus alphagastroenteritidis*) (primates). The presence of these viruses in primates is concerning as non-human primates are phylogenetically closely related to humans, which can facilitate interspecies viral transmission. These findings underscore the high diversity of mammalian viruses and the potential risk of human infection through cross-species transmission during the close interactions with wildlife in the wild meat supply chain.

## INTRODUCTION

Most human infectious diseases originate from animals, including domestic and wild animals (Jones et al., 2008; L. H. Taylor et al., 2001). Especially in tropical regions, wild mammals are an important reservoir of zoonotic pathogens (Wolfe et al., 2007). One significant path for the transmission of these pathogens from wildlife to humans is through activities involved in the supply chain of wild meat, defined here as meat from non-domesticated mammals — also commonly referred to as "bushmeat" in tropical regions (Milbank & Vira, 2022). These activities include intimate human-wildlife interactions, such as hunting, butchering, selling, cooking, and consumption, which facilitate the transmission of zoonotic pathogens to humans (Huong et al., 2020; Lucas et al., 2022; van Vliet et al., 2022). Between 1940 and 2021, most zoonotic spillover events associated with the wild meat supply chain occurred in Africa, where viruses were the most frequently reported pathogens (Milbank & Vira, 2022). Over a quarter of mammalian species involved in the wildlife trade, including the trade of wild meat, harbour 75% of known zoonotic viruses (Jones et al., 2008; Shivaprakash et al., 2021; Taylor et al., 2001). Notable examples of viral disease outbreaks that have been linked to contact with wild meat are Mpox (Okareh & Morakinyo, 2018), Ebola disease (Judson et al., 2016), SARS, and COVID-19 (Bell et al., 2004; Lytras et al., 2021).

In many parts of the Afrotropics, hunted wildlife traditionally serves as a source of protein, micronutrients, and income for many rural communities (Fa et al., 2002; Wilkie et al., 2016). Rapid population growth and urbanisation, the availability of firearms, and the increased rainforest accessibility through the expansion of road networks and logging, have led to unsustainable levels of commercial hunting of wildlife species (Cawthorn & Hoffman, 2015; Kleinschroth et al., 2019). Thus, in addition to public health concerns, the hunting of wildlife for their meat poses a significant threat to the conservation of tropical animals, driving many species towards extinction and potentially disrupting ecosystem services and functioning (Cawthorn & Hoffman, 2015; Fa & Brown, 2009; Ripple et al., 2016). The urban demand is mainly driven by the taste of wild meat, cultural connotations, and the perception that wild meat is more ‘pure’ and healthier than domestic animals, and many people are willing to pay higher prices than for domestic meat (Bair-Brake et al., 2014; Cawthorn & Hoffman, 2015; Lucas et al., 2022). As a result, wild meat consumption has well expanded beyond the geographical area where the wild meat was harvested and the high demand from international diaspora leads to a further intercontinental spread of these exotic animal products and the potential pathogens that they may carry (Bair-Brake et al., 2014; Chaber et al., 2023; Falk et al., 2013; Morrison-Lanjouw et al., 2023). In Europe, high quantities of unregulated wild meat are imported. For example, in Belgium, approximately 3.9 tonnes of wild meat enter illegally each month via Brussels Zaventem airport, primarily originating from sub-Saharan Africa, with the Democratic Republic of the Congo (DRC) being the main source (Chaber et al., 2023; Gombeer et al., 2021).

Our understanding of the wildlife virome in African mammals and their meat is limited and skewed towards species that are feasible to sample in the wild and those with a history of reported zoonotic disease emergence (Albery et al., 2020; Chen et al., 2023; Johnson et al., 2020; Shaw et al., 2020). These include small mammal species with large populations, such as common gregarious bats, rodents, and shrews. Consequently, this knowledge gap restricts our understanding of the origins of many infectious diseases and the potential risk of viral emergence in humans. To bridge this gap, systematic surveys across a broad range of wildlife taxa are necessary (Holmes et al., 2024). In this regard, conducting viral surveillance of specimens intended for the wild meat trade offers valuable opportunities to screen a wide variety of mammal species that are often challenging to sample due to their rarity, vulnerability, or protected status (Fa et al., 2002; Ripple et al., 2016; Wacharapluesadee et al., 2021; Wilkie et al., 2016) and to provide a better understanding of the viruses circulating in wildlife that may come into direct contact with humans via the wild meat supply chain.

Alongside the significant bias in host surveillance, our understanding of the wildlife virome is disproportionately focused on specific viruses. Wildlife pathogen screening typically focuses on known viruses with reported public health or economic consequences and relies on short genetic sequences instead of complete viral genomes (Wacharapluesadee et al., 2021). However, to uncover the true diversity of viruses in wildlife samples, it is imperative to employ unbiased viral detection techniques that target whole genomes, regardless of prior knowledge of the host-virus associations. Such unbiased detection methods use recent advancements in next-generation sequencing technologies, providing high-quality genomic sequencing data, even for challenging samples with degraded DNA or RNA, such as processed (i.e., smoked/cooked) wild meat (He et al., 2022; Katani et al., 2021; Temmam et al., 2017; Zhang et al., 2019).

In the present study, we use viral metagenomics to analyse a collection of tissue and swab samples from mammal specimens intended for the wild meat trade in several regions of the DRC and Brussels, Belgium. Our goals are to detect and characterise the viral diversity in these wildlife species and to assess the potential health risks associated with activities related to the wild meat supply chain.

## MATERIAL AND METHODS

### Sample collection and mammal species identification

Samples were collected from mammals that were intended for the meat trade (i) at African grocery stores in the Matongé neighbourhood in Brussels, Belgium, in 2017 and 2018 (ten samples) (Gombeer et al., 2021), (ii) from freshly killed animals in Inkanamongo in the Tshuapa province, DRC, in 2021 (53 samples) (van Vredendaal et al., 2024), and (iii) from archived wild meat stored at the collection of the Royal Belgian Institute of Natural Sciences (RBINS) and University of Antwerp (UAntwerp) (36 samples). The archived wild meat was collected at several rural and urban markets in the Tshopo, Ituri, and Bas-Uélé provinces (DRC) between 2010 and 2015 (**Figure 1**; **Supplementary Table S1**).

**Figure 1.**
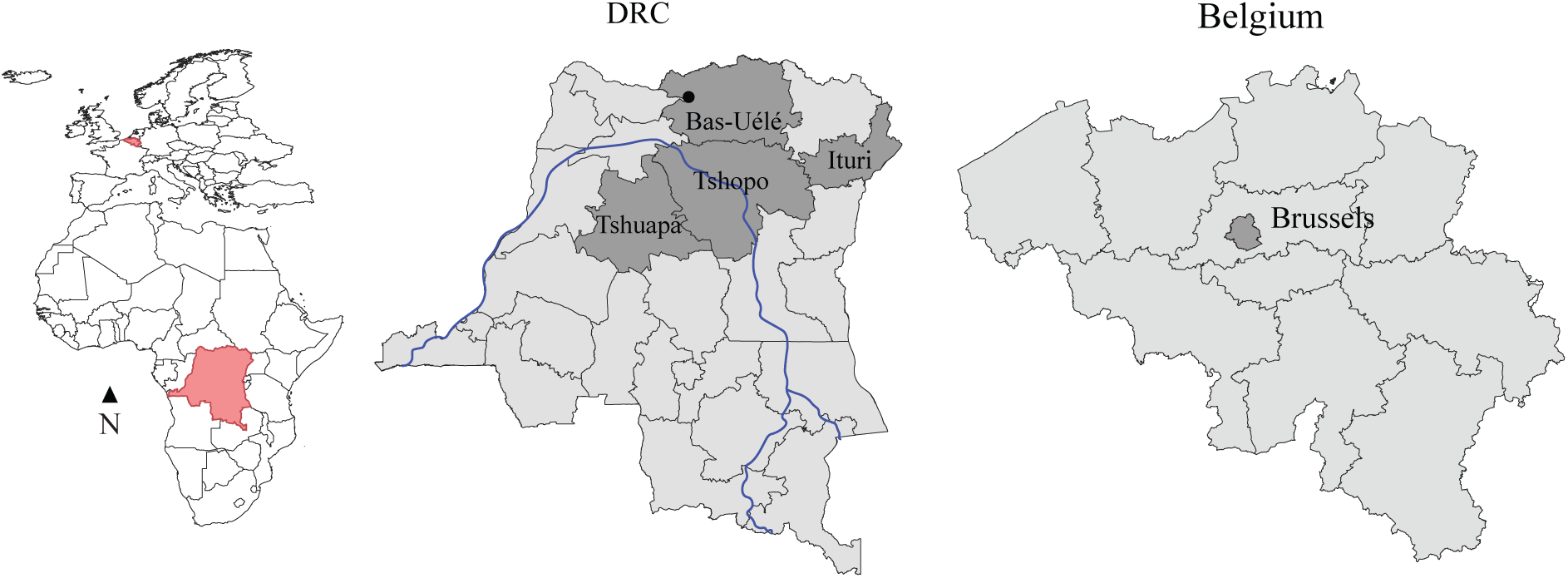
Map showing the provinces in the DRC and Belgium where samples were collected. On the left, a map of mainland Africa and Europe, with the DRC and Belgium marked in red. The provinces where samples were collected are shaded in dark grey in each country. The main course of the Congo River in blue.

Carcasses from the African grocery stores in Brussels were obtained as described in Gombeer et al. (2021) in November − December 2017 and May 2018, and frozen at -20°C. The carcasses were partially unfrozen to collect three pieces of muscle and/or bone marrow, which were stored at -20°C until further processing. The samples collected in Inkanamongo consisted of an oral, nasal, rectal and urogenital swab per carcass, which were kept in DNA/RNA Shield in the field for three to five weeks and later transferred to -80°C for long-term storage. The samples from the archived collection of RBINS and UAntwerp consisted of samples of various tissues from both fresh and processed wild meat, which have been stored at room temperature in ethanol or frozen at -20°C in RNAlater (**Supplementary Table S1**). DNA was extracted from the samples using the QIAamp® DNA Micro kit or the DNeasy Blood & Tissue kit (spin column or plate extraction kits, Qiagen), following the manufacturer’s instructions. Molecular identification of the mammal species was performed by using a multiplex DNA-barcoding approach following Gaubert et al. (2015), which targets four mitochondrial genes: the cytochrome *b* gene (*cytb*), the cytochrome *c* oxidase subunit I gene (*COI*), and the ribosomal subunits 12S and 16S. The multiplex PCR products were sequenced using the Oxford Nanopore Technology on a Flongle flow cell. If a target sequence was not recovered, it was re-amplified in a singleplex PCR and Sanger sequenced by Macrogen Europe.

### RNA extraction

RNA from muscle and bone marrow samples collected in Brussels was extracted separately using the Nucleospin® RNA kit (Macherey-Nagel) following the manufacturer’s instructions, eluting with 60 μL RNase-free water. These extracts were pooled and concentrated to 35 µL using the RNeasy MinElute Cleanup Kit (Qiagen).

Prior to RNA extraction of the samples collected in the Tshuapa province, we pooled 70 µL of the DNA/RNA Shield medium of each of the four swabs. RNA extraction was performed with the QIAmp Viral RNA extraction kit (Qiagen) following the manufacturer’s instructions but substituting Qiagen’s columns for Zymo-Spin columns IIC (Zymo Research).

The RNA from samples collected from the archival collection was extracted using the Nucleospin® RNA kit (Machery-Nagel) following the manufacturer’s instructions, but without adding β-mercaptoethanol during the lysis step.

### PCR-based virus detection

As an initial viral screening, we conducted multiple PCR tests targeting viruses frequently associated with human epidemics, i.e., *Coronaviridae*, *Filoviridae* (e.g. Ebola virus, Marburg viruses), *Flaviviridae* (e.g. hepatitis C virus), *Paramyxoviridae* (e.g. canine distemper virus, measles virus, parainfluenza virus), simian immunodeficiency virus (SIV) and *Orthopoxvirus*, *Poxviridae* (e.g. monkeypox virus) (Nathanson, 2016). To screen for RNA viruses, RNA extracts were reverse-transcribed using Maxima Reverse Transcriptase (Thermofisher) following the manufacturer’s instructions and PCRs were carried out on the resulting cDNA. To screen for DNA viruses belonging to *Orthopoxvirus*, the DNA extracts were subjected to a quantitative real-time PCR (see **Supplementary Table S2** for information on primers and cycling conditions). PCR products were run on a 1.5% agarose gel and Sanger sequenced at the University of Antwerp VIB sequencing facility.

### Library preparation and next-generation sequencing (NGS)

For library construction, RNA extracts were pooled according to mammalian taxon and country of sampling with one to fourteen individuals per pool (**Supplementary Table S1**). For each pool, RNA was purified using the RNeasy MinElute Cleanup Kit (Qiagen) and quantified with the Qubit™ 2.0 Fluorometer (Life Technologies). Samples were sent to Genewiz (Azenta Life Sciences) for cDNA generation, library preparation, and sequencing using Strand-Specific RNA-Seq. The library preparation included an rRNA depletion step to remove host rRNA from the libraries, and, thereby, relatively increase the yield of viral RNA reads. Libraries were 150 bp paired-end sequences on an Illumina NovaSeq™ platform.

### Virus detection and genome assembly

The overall quality of the reads in each library was checked with FastQC v0.11.7 (Andrews, 2010). Subsequently, reads were paired with Trimmomatic v0.36 (Bolger et al., 2014) and (part of) the reads with a low-quality score were discarded, i.e., when the Phred-33 quality score within a window of four bases was below 15 and when the read was less than 50 bp long. To filter out reads originating from the host genome, trimmed reads were aligned against the corresponding host genome (or a phylogenetically closely related host genome in case the respective host genome was not available in the database of the National Center for Bioinformatic Information (NCBI) (https://www.ncbi.nlm.nih.gov/)) using the BWA-INDEX and BWA-MEM algorithm of BWA v0.7.17 (Li & Durbin, 2010) (**Supplementary Table S3**). The unmapped reads were *de novo* assembled into contigs using the rnaviralSPAdes assembler of SPAdes v3.15.5. Next, the resulting contigs were mapped against a vertebrate virus database using BLASTn (BLAST+ v2.13.0) with a word size of 16 and an E-value cut-off of 10^-10^. This vertebrate virus database was compiled from NCBI Virus (https://www.ncbi.nlm.nih.gov/labs/virus/) and includes all complete non-human vertebrate viral genomes (48,943 genomes) supplemented with all complete RefSeq human virus genomes (372 genomes). Contig hits with an alignment length of at least 100 bp were retained and blasted against the NCBI non-redundant nucleotide (nt) database (available as of May 8, 2024) using BLASTn to retain only contigs that matched to vertebrate virus sequences in the nt database. These contigs were then mapped to their respective blast hit reference genome in Geneious Prime v2024.0.3 at the highest sensitivity settings, iterating up to ten times. Quality-trimmed paired reads were mapped back to the consensus of the assembled contigs (majority consensus with the reference genome sequence value when there was no coverage) using Bowtie2 v2.4.4 with the local alignment mode ‘--very-sensitive-local’ to assess coverage (i.e., percentage of the reference sequence covered by reads) and depth (i.e., the average number of reads that detected a certain base in the genome), and the resulting consensus (majority consensus with N called when there is no coverage) was blasted against the NT database using BLASTn. The abundance of each virus species in the library was expressed as the number of mapped reads per million total (quality trimmed) reads (RPM). Following Cui et al. (2019, 2023), we considered a library to be false positive when the RPM was lower than 1. These ‘false positives’ were not included in further phylogenetic and recombinant analyses. We did, however, check whether the presence of these viral reads could have been caused by contamination from other viral positive libraries by mapping the viral reads against the viral consensus sequences of the ‘true positive’ pool for the respective viral family (if present) in Geneious Prime.

### Recombination and phylogenetic analyses

The virus sequences were assigned to a viral (sub)family based on the lineage information of the top hit in the BLASTn results (**Supplementary Table S3**). To verify this preliminary identification of the virus family and to further assign the virus to genus (and species), we performed recombination and phylogenetic analyses based on the RNA-dependent RNA polymerase (RdRp) gene sequence for RNA viruses, and the DNA polymerase gene sequence for DNA viruses as replication genes are the most conserved phylogenetic markers for viruses (Wolf et al., 2018; Zhang et al., 2019).

First, these genes were annotated in Geneious Prime using the annotated genomes of one virus exemplar per species in the respective family (selected based on the Virus Metadata Resource (VMR) from the International Committee on Taxonomy of Viruses (ICTV)) as a reference database and using a 50% identity threshold to transfer reference annotations. The annotated genes were further verified by identifying open reading frames (ORFs) using the Find ORFs tool in Geneious Prime. In case the replication gene was not assembled, the recombination and phylogenetic analyses were based on fragments of other genes: e.g., fragments of the envelope glycoprotein B (gB) and capsid gene were used for the analyses of the gammaherpesvirin strain that was detected in pool 45 (see Results section) (**Supplementary Table S4**).

Next, for each detected virus, we created two alignments: one that included a single sequence from each species in the respective family (the same database as for gene annotation), and another encompassing all viral sequences of the detected genus available in NCBI Virus to cover a broader phylogenetic diversity. However, for the genus *Orthohepadnavirus,* we only included specimens of species that clustered within the same clade as the detected virus in the family-level phylogeny (see Results section) to keep the dataset manageable.

At the family level, the alignment of the RdRp (or DNA polymerase) genes was performed using Translation Align in Geneious Prime at default settings. At the genus level, MAFFT v7.490 was used at default settings in Geneious Prime because not all RdRp fragments in the database were in the same translation frame. Also, short fragments, i.e. fragments shorter than 90% of the total expected length of the replication gene, were removed from the alignment. Subsequently, all positions with gaps were removed using TrimAl v1.2 (Capella-Gutiérrez et al., 2009). For retroviruses of the subfamily *Spumaretrovirinae* (family *Retroviridae*) (see Results section), we also performed recombination and phylogenetic analyses with a shorter RdRp sequence (271 bp after trimming and gap removal) to include a substantially larger amount of GenBank sequences into the analyses, i.e., 543 sequences instead of the 34 that were included with the longer RdRp fragment).

To test for the presence of recombination, we performed a PHI-test (Bruen et al., 2006) implemented in SplitsTree4 v4.19.2 (Huson, 1998) in combination with the recombination detection methods BootScan (window size 200) (Martin et al., 2005), Chimaera (variable window size) (Posada & Crandall, 2001), MaxChi (variable window size) (Posada & Crandall, 2001), RDP (window size 100 bp) (Martin & Rybicki, 2000), 3Seq (Lam et al., 2018), and SisScan (window size 200) (Gibbs et al., 2000) implemented in RDP4 v4.101 (Martin et al., 2015). In case the PHI-test was significant, and a recombination event was detected by at least 3 detection methods in RDP4, the recombinant region was excluded from the alignment prior to conducting the phylogenetic analyses.

To infer Maximum Likelihood (ML) trees, we used IQ-TREE v2.2.2.6 (Nguyen et al., 2015) with model selection based on the Bayesian Information Criterion (BIC) using ModelFinder (Kalyaanamoorthy et al., 2017) and 1,000 bootstrap replicates (**Supplementary Table S4**). Phylogenetic trees were visualised and midpoint-rooted with FigTree v1.4.4. Pairwise nucleotide *p*-distances (*p =* the proportion of nucleotide positions where the two compared sequences differ) were estimated using MEGA11 v11.0.13 (Tamura et al., 2021) using the gamma parameter and rates among sites as determined by IQ-TREE. Trees were visualised in Rstudio v2024.04.2+764 using the packages ‘ape’ v5.5 (Paradis et al., 2004) and ‘ggtree’ v3.0.2 (Yu, 2020).

## RESULTS

### Host identification

The complete collection of mammal samples comprised 99 samples. Molecular barcoding revealed the animals belonged to 28 species from 24 mammal genera and 17 families: Bovidae (n = 8) and Suidae (n = 2) from the order Artiodactyla; Cercopithecidae (n = 23), Galagidae (n = 1), Hominidae (n = 1), and Lorisidae (n = 1) from the order Primates; Herpestidae (n = 1) and Nandiniidae (n = 3) from the order Carnivora; Hystricidae (n = 27), Muridae (n = 1), Nesomyidae (n = 5), Sciuridae (n = 6), and Thryonomyidae (n = 4) from the order Rodentia; Macroscelididae (n = 13) from the order Macroscelidea; Manidae (n = 1) from the order Pholidota; Procaviidae (n = 1) from the order Hyracoidea; and Tenrecidae (n = 1) from the order Afrosoricida. For some mammals, species identification was inconclusive: the procavid in pool 11 was identified as a species of *Dendrohyrax*; the lorisid in pool 28 was identified as a species of Perodicticinae; one of the rodents in pool 31 was identified as a species of *Tryonomys*; bovids in pool 32 were identified as either domestic cow, *Bos taurus*, or zebu, *B. indicus*; bovids in pool 33 and pool 42 were identified as Peter’s duiker, *Cephalophus callipygus*, Ogilby’s duiker, *C. ogilbyi*, or Weyns’s duiker, *C. weynsi* sp. 1 (pool 42) and sp. 2 (pool 33); the two suids in pool 35 were identified as either red river hog, *Potamochoerus porcus*, or bushpig, *P. larvatus*; and four cercopithecids in pool 48 were identified as either Wolf’s monkey, *Cercopithecus wolfi*, or Dent’s monkey, *C. denti*.

Samples were pooled into a total of 30 libraries for metagenomic sequencing (**Supplementary Table S1**).

### Virus detection and characterisation

None of the specimens tested positive in the PCR assays targeting viruses of *Coronaviridae*, *Filoviridae*, *Flaviviridae*, *Paramyxoviridae*, or *Orthopoxvirus*. However, all assays are known to miss some members of the targeted virus families or genus. One specimen of *Allenopithecus nigroviridis* hunted in Tshuapa province, DRC tested positive for SIV, which is included in pool 51 in the metagenomic screening (see further).

The number of reads per library in the metagenomic screening varied between 113 and 840 million reads with an average of 195 million reads. After trimming and host filtering, the number of reads remaining for *de novo* assembly and subsequent blasting of the contigs to the vertebrate virus database for viral discovery ranged between 0.2 and 152 million reads with an average of 46 thousand reads. In fourteen of the 30 libraries viral reads were detected, ranging from 2 reads to 3 million reads and an RPM ranging from 0.011 to 17,679.591 (**Supplementary Table S3**).

Despite the large number of viral reads detected, we focused on vertebrate-associated viruses with an RPM>1. Using this criterion, we detected viruses in eleven of the 30 pools. These included samples from the liver of a fresh carcass of a tree hyrax (pool 11) and fresh carcasses of *Funisciurus* (pool 30). We also found viruses in the muscle and/or bone marrow of De Brazza’s monkeys, *Cercopithecus neglectus* (pool 38), of which the meat was smoked. Additionally, we detected viruses in combined swabs (oral, nasal, rectal, and urogenital) from fresh carcasses of various species, including blue duiker, *Philantomba monticola* (pool 43), African palm civet, *Nandinia binotata* (pool 45), Allen’s swamp monkey, *Allenopithecus nigroviridis* (pools 47 and 51), Wolf’s monkey or Dent’s monkey (pool 48), red-tailed monkey, *Cercopithecus ascanius* (pools 49 and 50), and the sample pool including African brush-tailed porcupine, *Atherurus africanus*, and giant pouched rat, *Cricetomys* sp. 2 (sensu Olayemi et al. (2012)) (pool 53). The identified viruses belong to two DNA virus families (*Hepadnaviridae* and *Orthoherpesviridae*) and six RNA virus families (*Arteriviridae*, *Picobirnaviridae*, *Picornaviridae*, *Retroviridae* (subfamily *Spumaretrovirinae*), *Sedoreoviridae*, and *Spinareoviridae*) (**Table 1**).

**Table 1.**
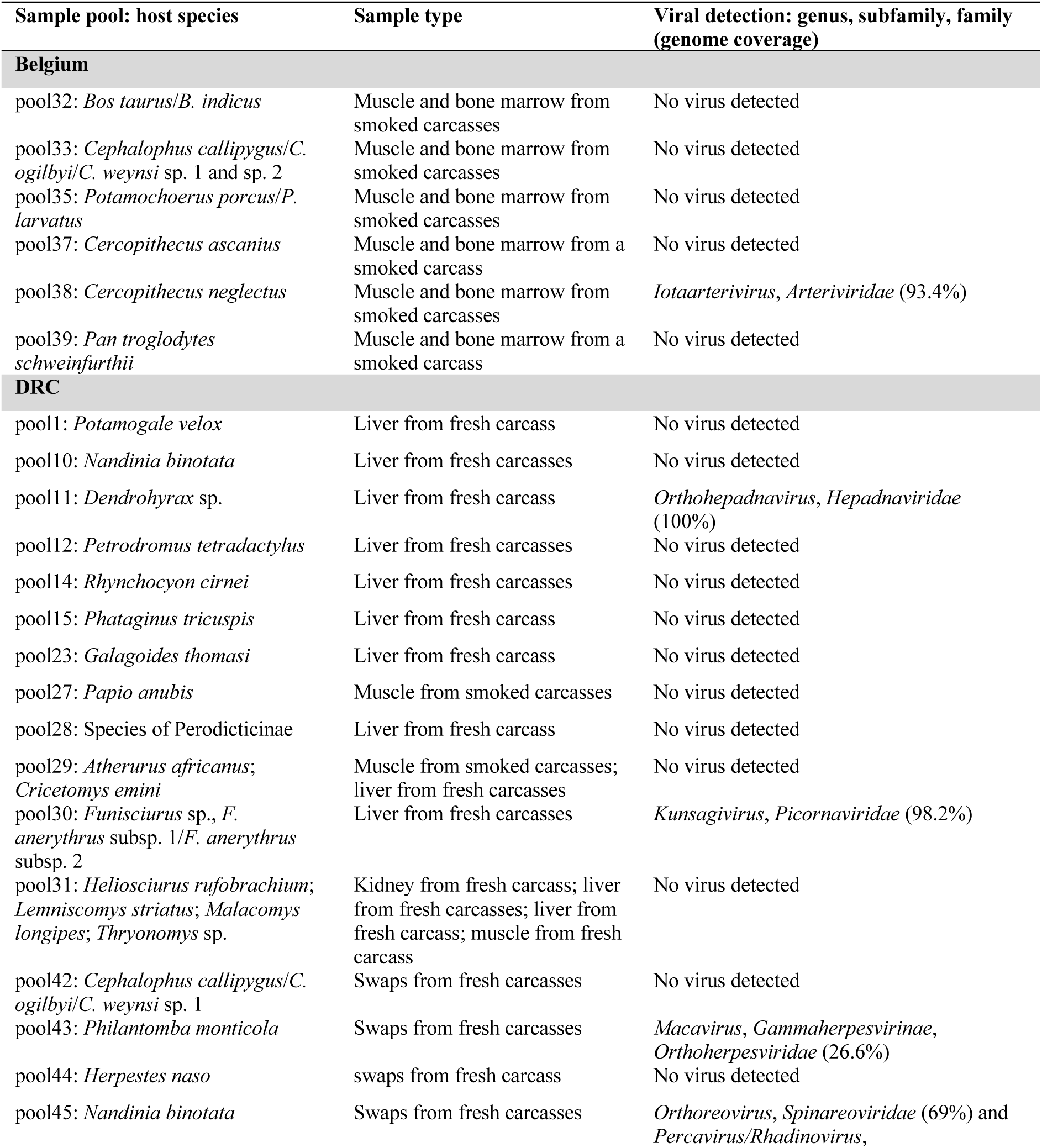
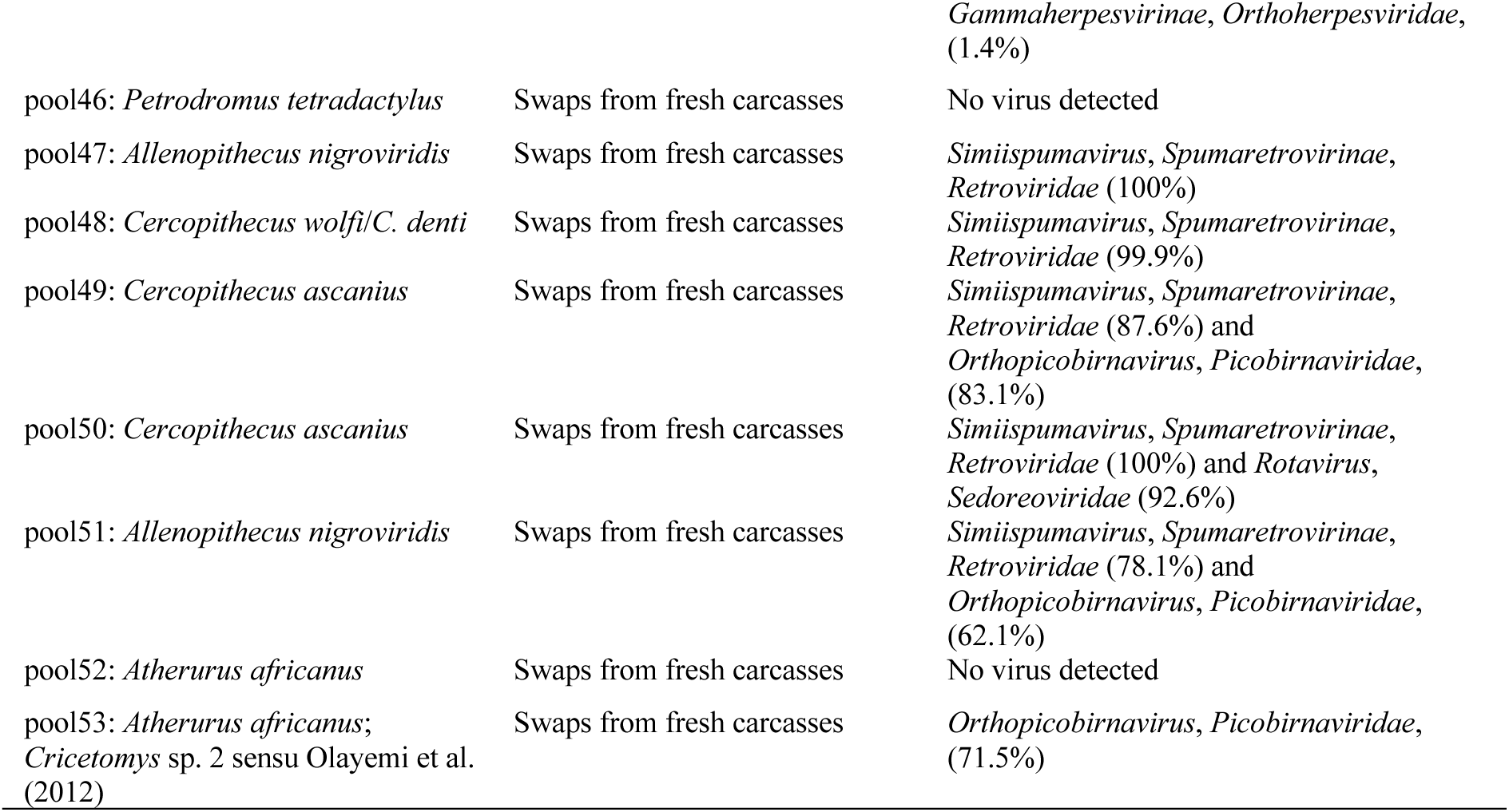
Viral detection in the read pools including information about the host species, sample type, and viral genus, subfamily (if applicable), family, and genome coverage.

For the detected viruses of which RPM < 1, we only found potential contamination in pool 14 and pool 38 of viral reads belonging to the family *Hepadnaviridae* from pool 11 as the viral reads (eighteen in pool 14, two in pool 38) mapped with 99.6% and 100% identity, respectively, against the consensus sequence belonging to *Hepadnaviridae* from pool 11. For the other pools with RPM < 1, viral reads had multiple SNPs with the consensus sequences of ‘true’ positive pools. Additional (deeper) sequencing and/or targeted PCR screening, and characterisation of the host genome could help determine whether these samples are truly negative and/or endogenous. This also applies to pools in which viral reads were detected but which were not tested for contamination as no confident positive pools were detected for the respective viral family. These include viral reads with top blast hits against the family *Flaviviridae* (porcine pegivirus) (pool 10), an unclassified virus (Ribovira sp.) (pool 14), the family *Astroviridae* (astrovirus sp.) and the family *Papillomaviridae* (Puda puda papillomavirus 1) (pool 42), unclassified viruses (Hudisavirus sp. and ticpantry virus 8) (pool 48), the family *Papillomaviridae* (Macaca mulatta papillomavirus 3) (pool 49), and the family *Flaviviridae* (simian pegivirus) (pool 51) (**Supplementary Table S3**). In pool 51, we detected a few viral reads (6) belonging to SIV (family *Retroviridae*, subfamily *Orthoretrovirinae*, genus *Lentivirus*, species *Lentivirus simimdef*). Although the RPM was below 1, one specimen included in this pool (EBO1480) tested positive for SIV using a conventional PCR assay, which confirms that this pool is indeed positive for this virus.

### Phylogenetic positioning of the detected viruses

We examined the phylogenetic relationships of the detected viral sequences at two levels: (i) at the family level to determine the virus genus (**Figure 2**), and (ii) at the genus level to offer a more detailed depiction of the evolutionary intrageneric relationships of the detected viruses (**Figure 3**; **Figure 4**).

**Figure 2.**
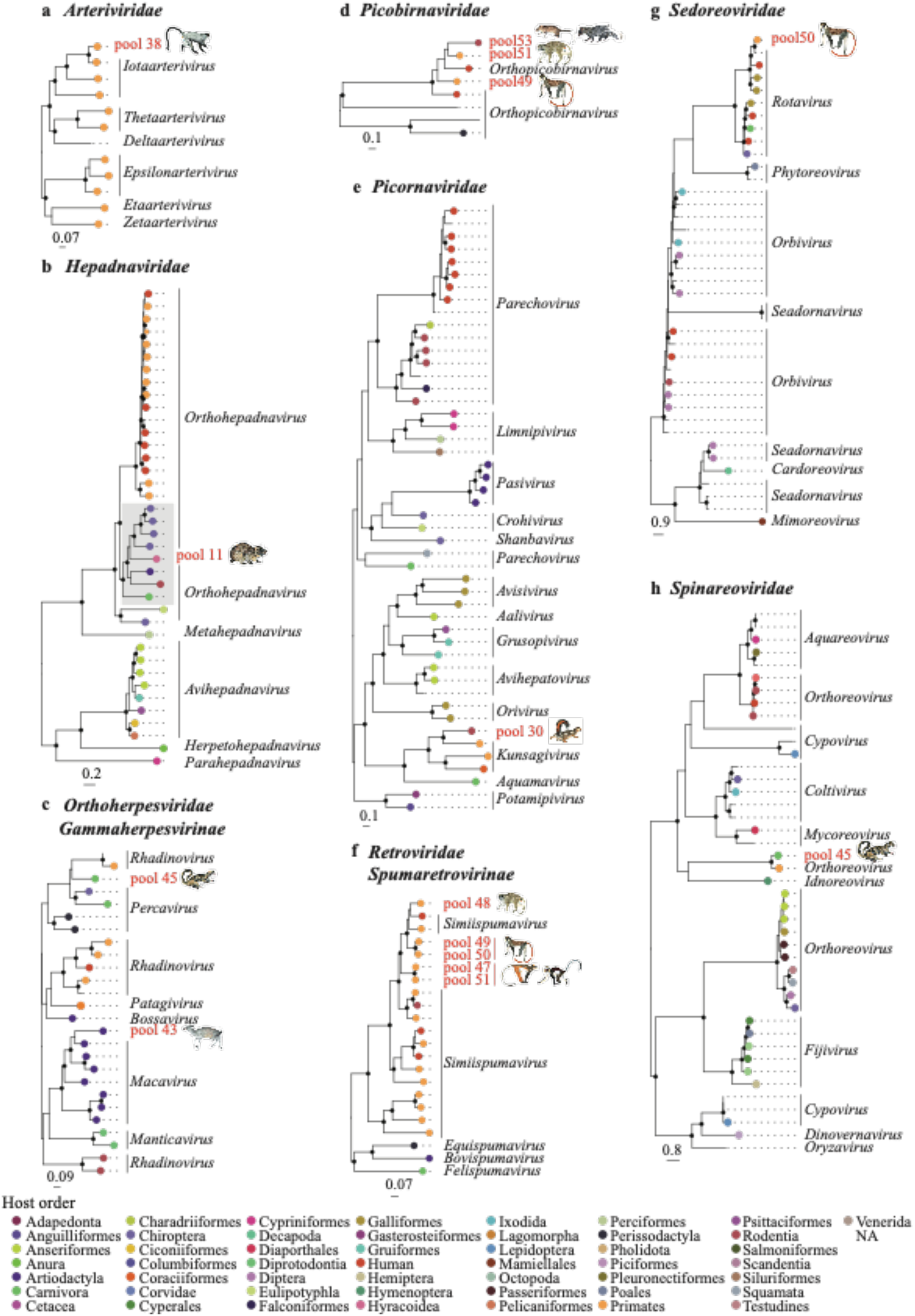
Phylogenetic relationships of members of *Arteriviridae* (a), *Hepadnaviridae* (b), *Gammaherpesvirinae* (*Orthoherpesviridae*) (c), *Picobirnaviridae* (d), *Picornaviridae* (e), *Spumaretrovirinae* (*Retroviridae*) (f), *Sedoreoviridae* (g), and *Spinareoviridae* (h) inferred from the RNA-dependent RNA polymerase gene (a, d-h), DNA polymerase gene (b), and capsid gene (c). Maximum likelihood phylogenetic trees with only well-supported nodes (bootstrap ≥ 85 %) indicated by black dots. Scale bar below each tree indicates the mean number of nucleotide substitutions per site. Tip points are coloured by the host order (humans indicated by a different colour than other primates). The tip label of the viral strain detected in the present study is in red and is labelled by the pool it was detected in. Mammal images of the hosts are from Kingdon (2015) and are depicted next to the respective pool. For pool 11, we depicted *Dendrohyrax arboreus*, for pool 30, we depicted *Funisciurus anerythrus*, for pool 53, we depicted *Cricetomys gambianus*.

**Figure 3.**
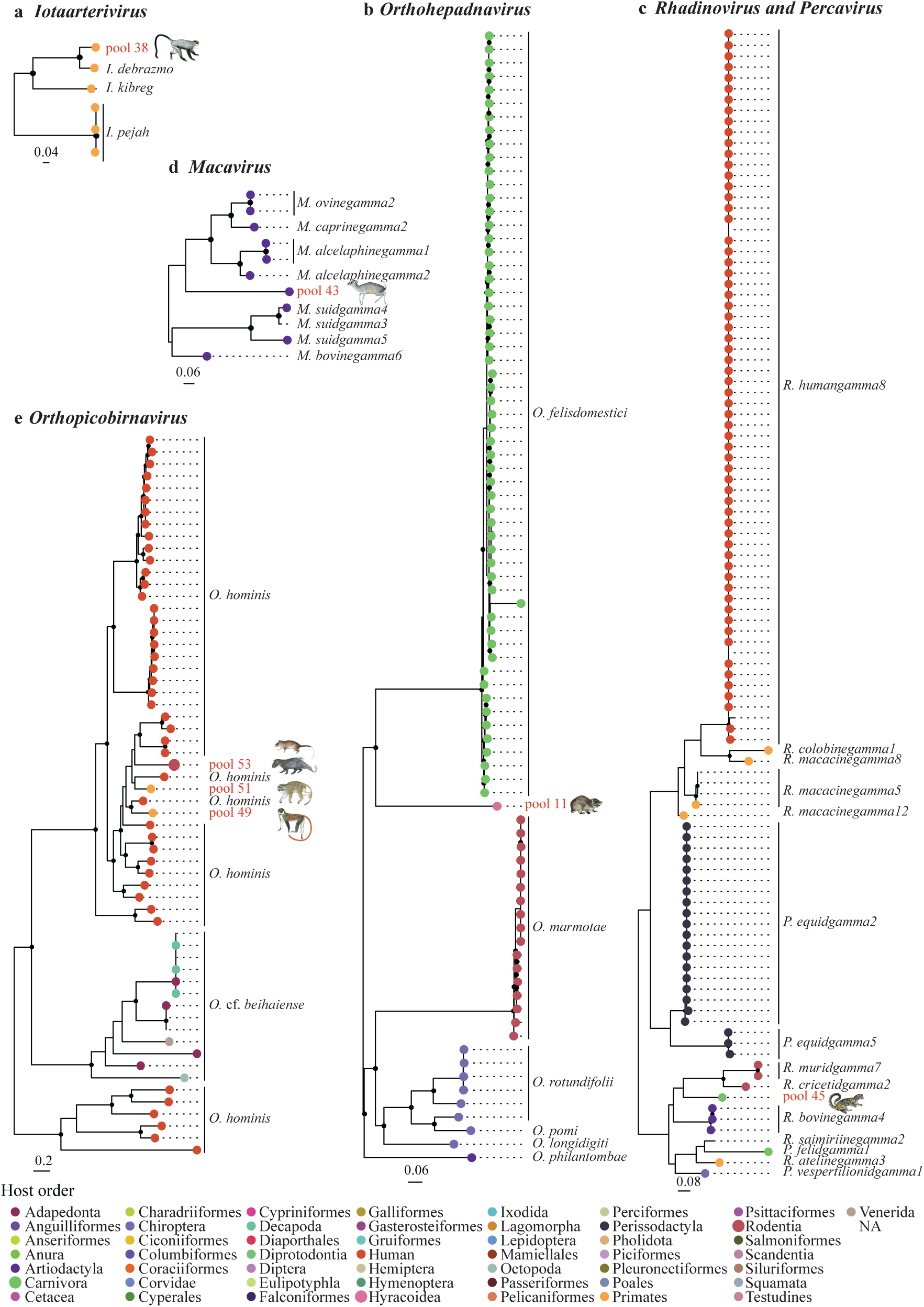
Phylogenetic relationships of members of *Iotaarterivirus* (a), *Orthohepadnavirus* (b), *Rhadinovirus* and *Percavirus* (c), *Macavirus* (d), and *Orthopicobirnavirus* (e) inferred from the RNA-dependent RNA polymerase gene (a, e), DNA polymerase gene (b, d), and capsid gene (c). Maximum likelihood phylogenetic trees with only well-supported nodes (bootstrap ≥ 85 %) indicated by black dots. Scale bar below each tree indicates the mean number of nucleotide substitutions per site. Tip points are coloured by the host order (humans indicated by a different colour than other primates). The tip label of the viral strain detected in the present study is in red and is labelled by the pool it was detected in. Mammal images of the hosts are from Kingdon (2015) and are depicted next to the respective pool. For pool 11, we depicted *Dendrohyrax arboreus*. For pool 53, we depicted *Cricetomys gambianus*.

**Figure 4.**
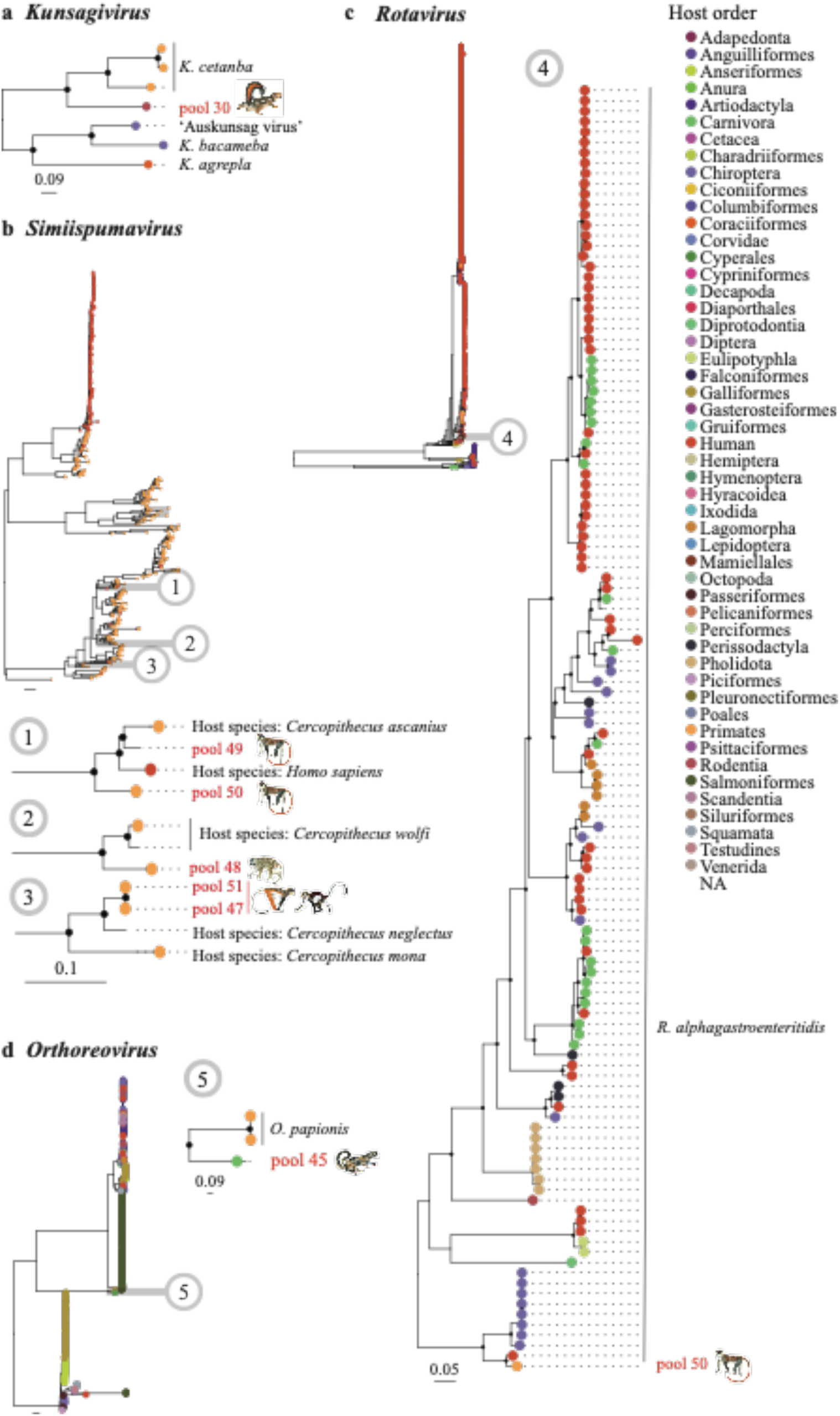
Phylogenetic relationships of members of *Kunsagivirus* (a), *Simiispumavirus* (b), *Rotavirus* (c), and *Orthoreovirus* (d) inferred from the RNA-dependent RNA polymerase gene (only a short fragment of 271 bp was used for (b)). Maximum likelihood phylogenetic trees with only well-supported nodes (bootstrap ≥ 85 %) indicated by black dots. Scale bar below each tree indicates the mean number of nucleotide substitutions per site. Tip points are coloured by the host order (humans indicated by a different colour than other primates). The tip label of the viral strain detected in the present study is in red and is labelled by the pool it was detected in. Mammal images of the hosts are from Kingdon (2015) and are depicted next to the respective pool. For pool 30, we depicted *Funisciurus anerythrus*.

The recombination analyses (Phi-test, BootScan, Chimaera, MaxChi, RDP, 3Seq, and SisScan) that were performed before each phylogenetic analysis (at both family and genus level) did not detect significant recombination events, i.e., the significance of the PHI-test and the detection of recombination events by at least 3 of the other detection methods (**Supplementary Table S5**). Note that for the recombination tests at the genus level for the rotavirus in pool 50, we only selected part of the sequences (i.e., the sequences within the clade depicted in **Figure 4-c** and **Supplementary Figure S10-b**) to maintain a manageable dataset.

### Arteriviridae (pool 38)

#### Iotaarterivirus

An arterivirus belonging to the genus *Iotaarterivirus* (family *Arteriviridae*, subfamily *Simarterivirinae*) was detected (genome coverage 93.4%) in the read pool from the muscle and/or bone tissue of two De Brazza’s monkeys, which were collected from smoked meat in a shop in Brussels (pool 38) in Gombeer et al., (2021) (**Table 1**; **Figure 2-a**; **Supplementary Table S6**; **Supplementary Figure S1-a**). It is closest related to DeBrazza’s monkey arterivirus (species *Iotaarterivirus debrazmo*) (GenBank accession number NC_026509) collected from the same primate species in Cameroon (*p*-distance = 0.144) (**Figure 3-a**; **Supplementary Table S7**; **Supplementary Figure S1-b**). A strain of *Iotaarterivirus kibreg* found in another species of *Cercopithecus*, *a* red-tailed monkey, further clusters with this group.

### Hepadnaviridae (pool 11)

#### Orthohepadnavirus

The strain belonging to the family *Hepadnaviridae* was detected (full genome coverage) in a liver sample from the fresh carcass of a tree hyrax from Tshopo (DRC) (pool 11). Phylogenetic analysis reveals that it clusters with members of the genus *Orthohepadnavirus* (**Table 1**; **Figure 2-b**; **Supplementary Table S6**; **Supplementary Figure S2-a**). This is the first time that this viral genus is detected in species of *Dendrohyrax*. In the intrageneric phylogenetic tree (**Figure 3-b**; **Supplementary Figure S2-b**), this strain groups with a clade containing members of domestic cat hepadnavirus (species *Orthohepadnavirus felisdomestici*) infecting domestic cats from Asia, Europe, North America, South America, and Australia, and with a clade containing members of roundleaf bat hepatitis B virus (species *Orthohepadnavirus rotundifolii*) infecting bats from Africa and Asia, long-fingered bat hepatitis B virus (species *Orthohepadnavirus longidigiti*) and pomona bat hepatitis B virus (species *Orthohepadnavirus pomi*) infecting bats from Asia, and Taï Forest hepatitis B virus (species *Orthohepadnavirus philantombae*) infecting an artiodactyl from Africa. Based on the pairwise nucleotide *p*-distances, the viral strain detected in the present study is closest related to roundleaf bat hepatitis B virus infecting bats of *Hipposideros* and *Rhinolophus* from Gabon (*p*-distance = 0.330 – 0.333) (**Supplementary Table S8-a**). As a full viral genome (3,188 bp) was recovered from this read pool, we additionally calculated the *p-*distances based on the full genome sequence (**Supplementary Table S8-b**), which shows the smallest *p*-distance between the viral strain detected in the present study and the strain of roundleaf bat hepatitis B virus infecting a bat of *Rhinolophus* from Gabon (*p-*distance = 0.327). According to the species demarcation criterion proposed for hepadnavirids (more than 20% nucleotide sequence divergence of the complete genomes) (*p*-distance > 0.20), the virus identified in this study can be considered as a new member of the genus *Orthohepadnavirus* which we tentatively name ‘Tree hyrax hepadnavirus’.

### Orthoherpesviridae (pool 43 and 45)

#### Percavirus or Rhadinovirus

A gammaherpesvirin strain (family *Orthoherpesviridae*, subfamily *Gammaherpesvirinae*) was detected in the pooled swab samples of a fresh carcass of an African palm civet (pool 45) from Inkanamongo, the first time that an orthoherpesvirid is detected in this mammal host (**Table 1**; **Figure 2-c**). Only a low percentage (1.4%) of the genome was covered (**Table 1**; **Supplementary Table S6**) with no coverage of the DNA polymerase gene. Therefore, we have used fragments of other genes, i.e., the capsid (**Figure 2-c**; **Figure 3-c**; **Supplementary Figure S3**) and gB gene (**Supplementary Figure S4**), to perform the phylogenetic analyses. Based on these analyses, the detected strain falls in the clade with members of the genera *Percavirus* and *Rhadinovirus*. Based on the capsid gene fragment, the lowest *p*-distance is found with members of vespertilionid gammaherpesvirus 1 (species *Percavirus vespertilionidgamma1*) (GenBank accession number KU220026) infecting a bat (*Myotis velifer*) from North America and saimiriine gammaherpesvirus 2 (species *Rhadinovirus saimiriinegamma2*) (GenBank accession number X64346) propagated in owl monkey kidney (*p*-distance = 0.283) (**Table S9-a**). Based on the gB gene fragment, the strain clusters together with members from the genus *Percavirus* and has the smallest *p*-distance to felid gammaherpesvirus 1 (species *Percavirus felidgamma1*) (GenBank accession numbers KT595939 and NC_028099) infecting cats (*Felis catus*) from North America (*p*-distance = 0.276) (**Supplementary Table S9-b**).

#### Macavirus

Another strain of gammaherpesvirus was detected (genome coverage 26.6%) in the pooled swab samples from fresh carcasses of two blue duikers (pool 43) from Inkanamongo (**Table 1**; **Figure 2-c**; **Supplementary Table S6**). This is the first time that an orthoherpesvirid is detected in this host. Based on phylogenetic analyses using fragments of the DNA polymerase (**Figure 3-d**; **Supplementary Figure S5**), capsid (**Supplementary Figure S3-a**), and gB (**Supplementary Figure S4-a**) genes, this strain clusters with members of the genus *Macavirus* (family *Orthoherpesviridae*, subfamily *Gammaherpesvirinae*) infecting artiodactyls from North America and Africa. Based on the pairwise *p*-distances of DNA polymerase gene fragment, the strain detected here was most similar to a strain of caprine gammaherpesvirus 2 (species *Macavirus caprinegamma2*) (GenBank accession number NC_043059) infecting goats (*p*-distance = 0.290) (**Supplementary Table S10**).

### *Picobirnaviridae* (pool 49, 51, and 53)

#### Orthopicobirnavirus

Three distinct strains belonging to the genus *Orthopicobirnavirus* (family *Picobirnaviridae*) were detected in swab samples of several mammal species: one in a sample pool from eight red-tailed monkeys (pool 49) (genome coverage 83.1%), one in the sample pool from two Allen’s swamp monkeys (pool 51) (genome coverage 62.1%), and one in the sample pool from thirteen northern giant pouched rat*s* and one African brush-tailed porcupine (pool 53) (genome coverage 71.5%) (**Table 1**; **Figure 2-d**; **Supplementary Table S6; Supplementary Figure S6-a**). All animals were sampled in Inkanamongo. According to the phylogenetic analyses using the RdRp gene fragment, all three strains cluster together with human picobirnavirus (species *Orthopicobirnavirus hominis*) from Asia and Europe (**Figure 3-e**; **Supplementary Figure S6-b**). Pool 49 and pool 51 have the smallest *p*-distances to a strain of human picobirnavirus from India (GenBank accession number AB517739) (*p*-distances = 0.236 and 0.252, respectively) (**Supplementary Table S11-a-b**), while pool 53 has the smallest *p*-distance with the strain from pool 51 (*p*-distance = 0.315) and pool 49 (*p*-distance = 0.338) (**Supplementary Table S11-c**).

### Picornaviridae (pool 30)

#### Kunsagivirus

In the pool of liver samples from fresh carcasses of five rope squirrels (*Fusciurus* spp.) from Tshopo (DRC) (pool 30) we detected a strain belonging to *Kunsagivirus* (genome coverage 98.2%), which is the first time that this viral genus is detected in this mammal genus (**Table 1**; **Figure 2-e**; **Figure 4-a**; **Supplementary Table S6**). According to the phylogenetic analyses, the detected strain clusters together with members of kunsagivirus C1 (or bakunsavirus) (species *Kunsagivirus cetanba*) (**Figure 4-a**; **Supplementary Figure S7**) with the closest sequence similarity to a bakunsavirus infecting *Papio cynocephalus* from Tanzania (GenBank accession number NC_034206; *p*-distance = 0.383) (**Supplementary Table S12**). As this is below the cut-off for species demarcation criterion proposed for the genus *Kunsagivirus* (<52% nucleotide difference within the 3CD gene), we assign the detected strain as belonging to the species *Kunsagivirus cetanba*.

### *Retroviridae* (pool 47, 48, 49, 50, and 51)

#### Simiispumavirus

Strains of simian foamy virus (SFV) (genus *Simiispumavirus*, family *Retroviridae*, subfamily *Spumaretrovirinae*) were detected in the swab samples from fresh carcasses of all primates collected in Inkanamongo: in pool 47 and 51 including Allen’s swamp monkeys (genome coverage 100% and 78.1%, respectively), pool 48 including Wolf’s or Dent’s monkey (genome coverage 99.9%), and pool 49 and 50 including red-tailed monkeys (genome coverage 87.6%, and 100%, respectively) (**Table 1**; **Figure 2-f**; **Figure 4-b**; **Supplementary Table S6**; **Supplementary Figure S8**). This is the first time that SFV is detected in Allen’s swamp monkey and Wolf’s or Dent’s monkey.

Based on the phylogenetic analyses of the short (271 bp) RdRp gene fragment (**Figure 4-b**; **Supplementary Figure S9**), strains of SFV detected in Allen’s swamp monkeys phylogenetically cluster with a strain infecting De Brazza’s monkey and Mona monkey, *Cercopithecys mona*, from the DRC (GenBank accession number MN325128 and MN325151, respectively). As species demarcation criteria include virus-host phylogeny, we tentatively call this strain ‘simian foamy virus Cercopithecus mona’. Strains of SFV detected in Wolf’s monkey or Dent’s monkey were genetically closely related to SFVs detected in Wolf’s monkey in the DRC (GenBank accession number JX157546 and JX157547). Finally, strains of SFV detected in the red-tailed monkey were genetically closely related to SFVs detected in the red-tailed monkey and humans from the DRC (GenBank accession number JX157541 and JX157543, respectively) (**Figure 4-b**; **Supplementary Figure S9**) (see **Supplementary Table S13** for *p*-distances). We tentatively call this strain ‘simian foamy virus Cercopithecus ascanius’.

### Sedoreoviridae (pool 50)

#### Rotavirus

A strain of the genus *Rotavirus* (family *Sedoreoviridae*) (genome coverage 92.6%) was detected in the pooled swab samples from fresh carcasses of two red-tailed monkeys collected in Inkanamongo (pool 50) (**Table 1**; **Figure 2-g**; **Figure 4-c**; **Supplementary Table S6**; **Supplementary Figure S10-a**). This is the first time that this viral genus is detected in red-tailed monkeys. Phylogenetically, this strain belongs to a clade that further includes rotavirus A (species *Rotavirus alphagastroenteritidis*) infecting a human from Kenya (GenBank accession number HM627553), giant-leaf-nosed bats, *Macronycteris gigas*, from Gabon (GenBank accession numbers MN528086, MN551587, MN477236, MN477225, MN528101, MN528075, and MN528116), and a Lander’s horseshoe bat, *Rhinolophus landeri*, from Nigeria (Genbank accession number OK087521) (**Figure 4-c**; **Supplementary Figure S10-b**). The smallest *p*-distance was found with the strain infecting a human from Kenya (GenBank accession number HM627553) (*p*-distance = 0.037) (**Supplementary Table S14**).

### Spinareoviridae (pool 45)

#### Orthoreovirus

In the read pool of the swab samples from a fresh carcass of an African palm civet from Inkanamongo (pool 45), we detected a strain belonging to the genus *Orthoreovirus* (family *Spinareoviridae*) (genome coverage 69.0%), which is the first time that this viral genus is detected in this host (**Table 1**; **Figure 2-h**; **Supplementary Table S6**). The detected strain phylogenetically clusters with baboon orthoreovirus (species *Orthoreovirus papionis*) infecting yellow baboons, *Papio cynocephalus* (Genbank accession numbers NC_015877 and HQ847903: identical sequences not subjected to final NCBI review) with *p*-distance = 0.357 (**Figure 4-d**; **Supplementary Figure S11**; **Supplementary Table S15**). According to the species demarcation criteria proposed for orthoreoviruses (i.e., more than 75% nucleotide sequence identity within a species), the strain detected in the present study represents a new orthoreovirus, which we will here tentatively refer to as ‘Civet orthoreovirus’ (Note that the *p*-distances calculated in the present study are only based on the RdRp gene sequence and that additional analyses based on other gene fragments could result in different nucleotide sequence identity levels).

## DISCUSSION

The trade of wild meat plays a pivotal role in the livelihood of millions of people from the Afrotropics (Fa et al., 2002; Fa & Brown, 2009; Wilkie et al., 2016). However, the elevated wildlife extraction, driven by increased demand from urbanized areas and international diaspora, not only threatens the conservation of African wildlife diversity but also facilitates the (international) spread of pathogens carried by these animals.

In this study, we performed a metagenomic viral survey of a total of 99 mammal samples, of which molecular barcoding showed that they belong to 27 wild African and one domesticated mammal species (**Supplementary Table S1**). All these mammals were traded for their meat in several regions in the DRC and the Matongé neighbourhood in Brussels, Belgium. As of 2024, the Convention on International Trade in Endangered Species of Wild Fauna and Flora (CITES) lists two of these mammal species, namely white-bellied pangolin, *Phataginus tricuspis*, and chimpanzee, *Pan troglodytes schweinfurthii*, in Appendix I (i.e., species threatened with extinction). These two species were sampled at markets in the Tshopo province (DRC). Nine other species, of which either the meat was sampled at markets or swabs were taken of fresh carcasses from hunted animals, are listed in Appendix II (i.e., not necessarily currently threatened with extinction but may become so unless trade is closely controlled), namely Ogilby’s duiker, blue duiker, Allen’s swamp monkey, red-tailed monkey, De Brazza’s monkey, Wolf’s monkey, olive baboon (*Papio anubis*), and angwantibos and pottos (Perodicticinae). The two species listed in Appendix I are also listed by the International Union for Conservation of Nature (IUCN) Red List of Threatened Species as endangered, and two of the species in Appendix II as near threatened (red-tailed monkey and Wolf’s monkey) (**Supplementary Table S16**). The presence of these species in our sample collection highlights the threat that wildlife hunting and trading pose to African wildlife diversity.

Despite the relatively limited sample size and low mammal diversity that was screened for viruses in the present study, we detected a high diversity of vertebrate viruses, i.e., fifteen strains belonging to eight viral families (*Arteriviridae*, *Hepadnaviridae*, *Orthoherpesviridae*, *Picobirnaviridae*, *Picornaviridae*, *Retroviridae*, *Sedoreoviridae*, and *Spinareoviridae*).

In the muscle and bone marrow samples of the smoked meat of two De Brazza’s monkeys sampled at the Matongé quarter in Brussels, we detected a simian arterivirus of the genus *Iotaarterivirus* (family *Arteriviridae*, subfamily *Simarterivirinae*) which phylogenetically clusters with DeBrazza’s monkey arterivirus (species *I. debrazmo*). Simarterivirins naturally infect various cercopithecoid primate species (i.e., Old World monkeys), causing subclinical infections. At present, they include six genera and eleven recognised monophyletic species (Bailey et al., 2016). Several of these viruses, such as simian hemorrhagic fever virus (SHFV) (genus *Deltaarterivirus*), simian hemorrhagic encephalitis virus (SHEV) (genus *Epsilonarterivirus*), and Pebjah virus (PBJV) (genus *Iotaarterivirus*) (Lauck et al., 2015), have been reported to infect and cause lethal simian hemorrhagic fever (SHF) in several species of captive Asian macaques after cross-species transmission from their natural host (Lauck et al., 2015; Vatter et al., 2015; Wahl-Jensen et al., 2016). Although simarterivirins are not known to infect humans, their zoonotic threat is considered high given (i) the fact that at least three distinct members of simarterivirins have already caused fatal infections in captive macaques after host-switching; (ii) the close phylogenetic relationship between humans and the natural cercopithecoid monkey host; and (iii) the immunological naivety of humans to arterivirids (Lauck et al., 2015; Vatter et al., 2015; Wahl-Jensen et al., 2016). In addition, Warren et al. (2022) found that SHFV can replicate in human cells *in vitro* and, therefore, may not require major adaptations to infect humans. Also, the high virus titres in the blood detected in naturally infected monkeys suggest that even minimal exposure to the blood could result in significant viral exposure to humans (Bailey et al., 2016). The detection of DeBrazza’s monkey arterivirus RNA in Brussels supports that potential zoonotic viruses are co-exported with their hosts’ meat outside of the country of origin. This underscores how the international wild meat trade extends the geographic reach of associated public health risks. However, the presence of viral RNA alone does not confirm whether the virus was infectious.

We detected an orthohepadnavirus (genus *Orthohepadnavirus*, family *Hepadnaviridae)* in the liver from a fresh carcass of a tree hyrax from Tshopo, which we have tentatively named ‘Tree hyrax hepadnavirus’.

Viruses belonging to the family *Hepadnaviridae* infect several mammal species. Hepatitis B virus (HBV) (species *Orthohepadnavirus hominoidei*) is a notorious example of an orthohepadnavirus that has caused a worldwide health problem, with chronic human infection leading to serious diseases, such as cirrhosis and liver cancer (Ott et al., 2012). Currently, ten strictly human-associated genotypes are recognised (Littlejohn et al., 2016), with some additional closely related strains infecting other apes, such as gorillas, chimpanzees, orangutans, and gibbons (Rasche et al., 2016; Starkman et al., 2003), and New World monkeys (de Carvalho Dominguez Souza et al., 2018; Lanford et al., 1998). Other viruses belonging to the genus *Orthohepadnavirus* have been found in Rodentia (species *O. marmotae* and *O. sciuri*) (Roth et al., 1985; Seeger et al., 1984), Artiodactyla (species *O. philantombae*) (Gogarten et al., 2019), Carnivora (species *O. felisdomestici*) (Aghazadeh et al., 2018), Eulipotyphla (species *O. soricisinensis*) (Nie et al., 2019), and Chiroptera (species *O. longidigiti*, *O. pomi*, *O. rotundifolii*, and *O. tabernarii*) (Drexler et al., 2013; He et al., 2013, 2015).

Hepadnavirids are considered to be highly host-specific and to have co-diverged with their host. However, several recombination and cross-species transmission events between human and other primate orthohepadnaviruses suggest a geographic rather than host-specific distribution of primate orthohepadnavirus variants, highlighting the importance of recent cross-species transmissions in the ecology and evolution of orthohepadnaviruses and their zoonotic potential (Littlejohn et al., 2016; Starkman et al., 2003; Takahashi et al., 2000). Similarly to primate orthohepadnaviruses, those infecting African bats (*Hipposideros* and *Rhinolophus*) and Asian grey shrews cluster together according to their geographic origin rather than host species (Drexler et al., 2013; Nie et al., 2019), further corroborating the importance of local cross-species transmissions in the evolution of orthohepadnaviruses. The orthohepadnavirus detected in the present study is phylogenetically closely related to an orthohepadnavirus infecting African bats, suggesting that past cross-species transmissions might not only have been limited to the host genus and family level but could also extend across mammalian orders (here Chiroptera and Hyracoidea), as suggested by Nie et al. (2019).

This study is the first to report a strain of *Orthohepadnavirus* in a species of *Dendrohyrax*, thereby expanding the host range of this viral genus. In addition, this study is only the second to report on the virome of species of Hyracoidea (one other study reports a simplexvirus (family *Orthoherpesviridae*, subfamily *Alphaherpesvirinae*, genus *Simplexvirus*), infecting rock hyraxes, *Procavia capensis* in captivity (Galeota et al., 2009)). These results underscore the importance of continuous surveillance and monitoring of wild, understudied mammal species.

Two gammaherpesvirins (family *Orthoherpesviridae*, subfamily *Gammaherpesvirinae*) were detected in the swab samples from fresh carcasses of the African palm civet and blue duiker from Inkanamongo. Orthoherpesvirids infect a wide array of vertebrate hosts, including mammals, birds, and reptiles, usually establishing a life-long, latent infection. Viruses belonging to this family are highly host-specific and generally show a long-standing co-evolution with their host, only causing severe disease in the foetus and very young or immunocompromised individuals (Ehlers et al., 2008; McGeoch et al., 2006). Despite this narrow host range, orthoherpesvirids have been reported to spill over to new (distantly related) hosts, causing serious disease and death, and raising the question of their zoonotic potential (Ehlers et al., 2008; Tischer & Osterrieder, 2010). Within the subfamily *Gammaherpesvirinae*, seven genera are currently recognised by the ICTV, of which members infect a wide array of mammal orders: a member of the genus *Bossavirus* infecting an aquatic artiodactyl (dolphin), members of the genus *Lymphocryptovirus* infecting primates, members of the genus *Macavirus* infecting artiodactyls, members of the genus *Manticavirus* infecting diprotodont marsupials (koalas and wombats), a member of the genus *Patagivirus* infecting bats, members of the genus *Percavirus* infecting perissodactyls and carnivores, and members of the genus *Rhadinovirus* infecting primates, rodents, and artiodactyls. The strain that is detected in the African palm civet phylogenetically clusters together with the members of *Percavirus* and *Rhadinovirus*. Given the low percentage of genome coverage and depth (**Supplementary Table S6**), we were not able to unequivocally assign this virus strain to genus level based on its phylogeny. However, given the high host specificity of gammaherpesvirins, we hypothesise that it concerns a member of *Percavirus* as this is the only genus within *Gammaherpesvirinae* of which species have so far been reported to infect carnivores. The strain detected in the blue duiker phylogenetically clusters with members of *Macavirus*, a genus within *Gammaherpesvirinae* of which members are known to infect artiodactylids.

We detected viruses of the genus *Orthopicobirnavirus* (family *Picobirnaviridae*) in the swab samples of fresh carcasses of several mammal species from Inkanamongo: in the red-tailed monkey, Allen’s swamp monkey, and in pooled samples of northern giant pouched rat and an African brush-tailed porcupine. All detected strains phylogenetically cluster with *Orthopicobirnavius hominis* found in diarrhoetic humans from Asia and Europe (**Supplementary Figure S6**) (Ganesh et al., 2011; van Leeuwen et al., 2010). Picobirnavirids have been linked to diarrhoea and gastroenteritis in terrestrial mammals and birds. However, the pathogenicity of these viruses is still under debate (Delmas et al., 2019). Recent research put forward considerable evidence that picobirnavirids may in fact represent a family of bacteriophages and/or mycophages, e.g., the majority of viruses has been detected in stool samples; picobirnavirids have not yet successfully been cultured in eukaryotic cell lines nor isolated from animal tissue samples; picobirnavirids are detected in variable non-animal sources (e.g., wastewater, farmland and forest soils, sewage, permafrost); the closest relative to the family *Picobirnaviridae* is a protist-infecting member of the family *Partitiviridae*; there is an apparent lack of phylogenetic congruence between the vertebrate host and virus; and there are bacterial genetic motives present in the viral genome (Sadiq et al., 2024; Wang, 2022; and references herein).

At present, only one genus, *Orthopicobirnavirus*, and three species (*O. hominis*, *O. equi*, and *O. beihaiense*) within the family *Picobirnaviridae* are formally ratified by the ICTV: the two former infecting vertebrates, the latter infecting invertebrates. However, Sadiq et al. (2024) proposes a reclassification based on the seven phylogenetic clusters they have found in an extensive dataset of picobirnavirids infecting mammals, birds, fish, reptiles, invertebrates, environmental samples and microbial communities: four genera comprising predominantly animal-associated picobirnaviruses (‘Alphapicobirnavirus’ (including *O. equi*); ‘Betapicobirnavirus’ (including *O. hominis*), ‘Epsilonpicobirnavirus’, and ‘Zetapicobirnavirus’) and three genera which are mainly detected in environmental samples and microbial communities (‘Gammapicobirnavirus’ (including *O. beihaiense*), ‘Deltapicobirnavirus’, and ‘Etapicobirnavirus’).

Although the association of picobirnavirids with vertebrate hosts or rather their microbiota remains unclear, extensive host switching of the virus (or its microbiotic host), including transmissions between humans and non-human animals, has been documented within animal-dominating picobirnavirids (Sadiq et al., 2024; Vanderhoeven et al., 2024). The potential spillover of these viruses from animals to humans is also supported by our findings, showing a high genetic similarity between picobirnavirids in primates and rodents, and those infecting humans. Given the potential link of these viruses to human disease, it is crucial to identify the true host(s) of picobirnavirids. This could be achieved by determining whether animal cells, bacteria, or fungi can support picobirnavirid infection and propagation, or by using immunofluorescence to investigate the intracellular presence of picobirnavirid proteins and RNA in tissue biopsies and enteric microbiomes of potential animal hosts (Wang, 2022).

Of note is that we did not detect any viral reads mapping to the genome segment encoding the capsid (picobirnavirids are characterised by a genome consisting of two double-stranded RNA segments), which suggests its absence. However, although several unsegmented picobirnavirus genomes have been reported recently (Giannitti et al., 2015; Luo et al., 2018; Ullah et al., 2022), the apparent absence of the segment encoding the capsid could also be ascribed to the inability to detect it due to its high sequence diversity (Wang, 2022).

A strain of kunsagivirus (family *Picornaviridae*, subfamily *Paavivirinae*, genus *Kunsagivirus*) was detected in the liver from fresh carcasses of rope squirrels belonging to *Funisciurus* from Tshopo, which phylogenetically clusters with kunsagivirus C1 (bakunsavirus) (species *K. cetanba*) infecting yellow baboons from Tanzania. Currently, three species within the genus *Kunsagivirus* are recognised: members of *K. agrepla* infecting birds (Boros et al., 2013), members of *K. bacameba* infecting bats (Van Brussel, 2023; Yinda et al., 2017), and members of *K. cetanba* infecting primates (Buechler et al., 2017; Kuhn et al., 2020). To the best of our knowledge, our study is the first one to report the presence of kunsagiviruses in rodents, thereby expanding the host range of this genus. Kunsagiviruses are currently not linked to any diseases, although further research is needed to investigate this.

In the pooled swab samples of all cercopithecids (Allen’s swamp monkey, red-tailed monkey, Wolf’s monkey or Dent’s monkey) from Inkanamongo, we detected strains of simian foamy virus (SFV) (family *Retroviridae*, subfamily *Spumaretrovirinae*, genus *Simiispumavirus*). The viruses detected in Allen’s swamp monkey phylogenetically cluster with SFV infecting De Brazza’s monkey and Mona monkey from the DRC. The SFV that is detected in Wolf’s monkey or Dent’s monkey clusters with other strains from the same hosts from the DRC. The two viruses detected in red-tailed monkeys fall in the same clade as the SFV detected in a red-tailed monkey and a human from the DRC. The fact that all sample pools of cercopithecids from Inkanamongo were positive could point to a high prevalence of simian foamy viruses in these non-human primates (NHPs). This is in line with the findings of Mouinga-Ondémé and Kazanji (2013) who found a high prevalence of these viruses in NHPs and primate meat from Gabon.

Simian foamy viruses naturally infect numerous NHPs, causing persistent, latent infection. Phylogenetic analyses show a strong correlation between the host and virus phylogeny, supporting the long-term co-evolution of SFVs with their natural host (Katzourakis et al., 2014; Khan et al., 2018; Switzer et al., 2005). This is also corroborated by the findings in the present study, where the detected SFVs phylogenetically cluster according to the host species (**Supplementary Figure S9**).

Although NHPs have their own host-specific SFVs resulting from this codivergence, they remain susceptible to SFVs from other primate hosts, as evidenced by cross-species transmissions and several reported zoonotic spillovers to humans (Katzourakis et al., 2014; Leendertz et al., 2008; Switzer et al., 2005). Simian foamy viruses replicate in the oral mucosa of their host and multiple (persistent) infections have been reported in humans from Africa, mostly resulting from bites inflicted during hunting (Betsem et al., 2011; Buseyne et al., 2018; Rua et al., 2012; Switzer et al., 2012). Zoonotic SFV infections are assumed to be nonpathogenic. However, recent research has reported haematological abnormalities and mild to moderate anaemia in humans from Cameroon infected by simian foamy virus Gorilla gorilla gorilla (species *Simiispumavirus gorgorgor*) (Buseyne et al., 2018).

The strains of SFV that we detected in the red-tailed monkey are genetically similar to a strain found in humans from the DRC by Switzer et al. (2012) (GenBank accession number JX157543). Interestingly, one of these strains (the one detected in pool 49) was genetically more similar to the human-infecting strain than the SFV strains that were isolated from red-tailed monkeys in the study of Switzer et al. (2012). This finding corroborates the zoonotic potential of SFVs in general and emphasises the potential risk for spillover of SFVs from red-tailed monkeys.

Up to date no secondary transmissions of SFVs between humans have been reported, though data on person-to-person spread is limited to a few studies that follow up a small number of people only for a short period of time (Switzer et al., 2012). Increased pathogenicity following cross-species transmission from NHPs and secondary transmission are of great concern given that other retrovirids, such as human immunodeficiency virus 1 and human immunodeficiency virus 2 (HIV) (family *Retroviridae*, subfamily *Orthoretrovirinae*, genus *Lentivirus*, species *L. humimdef1* and *L. humimdef2*, respectively) and human T-lymphotropic virus 1 (HTLV-1) (family *Retroviridae*, subfamily *Orthoretrovirinae*, genus *Deltaretrovirus*), also originated from NHPs and have caused serious pandemics (Peeters et al., 2002; Wolfe et al., 2005). This, together with the multiple zoonotic spillovers that have been recorded for SFV, highlight the need to determine exposure risk and evaluate secondary transmissibility and pathogenicity of SFVs in humans and their natural hosts (Switzer et al., 2012).

A strain of rotavirus A (RVA) (family *Sedoreoviridae*, genus *Rotavirus*, species *Rotavirus alphagastroenteritidis*) was detected in the swab samples of red-tailed monkeys from Inkanamongo. Rotaviruses cause diarrhoea-associated morbidity in many bird and mammal species (including humans) and are transmitted by a faecal-oral route. Especially strains of RVA are of great concern for public health, being the global leading cause of diarrheal disease and deaths among children younger than 5 years, with the highest mortality in sub-Saharan Africa (Troeger et al., 2018).

Rotaviruses are characterised by a linear dsRNA genome consisting of eleven segments encoding six structural (VP1, VP2, VP3, VP4, VP6, and VP7) and five non-structural (NSP1, NSP2, NSP3, NSP4, and NSP5) proteins. They show great intraspecific genetic diversity, which can be attributed to genome segment reassortment following co-infection of the same host cell by different strains, creating unique genotype constellations. Indeed, there are numerous examples in literature of genotype reassortments, often in combination with transmission between various mammalian species, including humans (Cook, 2004; Díaz Alarcón et al., 2022; Ghosh et al., 2011; Kia et al., 2021; Matthijnssens et al., 2010; Simsek et al., 2021).

Based on the phylogeny of the RdRp (VP1) gene, the strain that was detected in the present study clusters with a rare strain of RVA (strain B10) infecting an infant from Kenya (Ghosh et al., 2011) and strains found in Lander’s horseshoe bat from Nigeria (Kia et al., 2021) and giant roundleaf bats from Gabon (Simsek et al., 2021).

Genomic analysis of the RVA strain infecting the infant from Kenya (strain B10) by Gosh et al. (2011) suggested a simian origin (simian strain SA11) of eight genome segments (VP2, VP3, VP4, VP7, NSP1, NSP2, NSP3, and NSP5) while the origin of the VP1, VP6, and NSP4 genes could not be determined as they did not cluster with any known genotypes. Then, in 2021, Simsek et al. (2021) found high degrees of nucleotide similarity between bat RVAs from Gabon and Kenya and strain B10 for the VP1, VP6, and NSP4 genes as well as for VP2-4, NSP1, NSP3, and NSP5 genes, suggesting bats as the major host of this human strain. A bat origin was further corroborated by the findings of Kia et al. (2021) who found high nucleotide similarity between the VP1 genes of a strain of RVA from a bat from Nigeria and strain B10. However, in the present study, *p-*distances calculated based on the VP1 gene show higher nucleotide similarity between the strain detected in red-tailed monkey and strain B10 (*p*-distance = 0.037) compared to those between bats from Gabon or Nigeria and B10 (highest *p-* distance = 0.11). This result favours the hypothesis of a simian origin proposed by Ghosh et al. (2011). Additional phylogenetic analyses based on the other segments of the RVA genome (VP2-4, VP6-7, NSP1-5) (**Supplementary Table S17**) reveal the highest nucleotide similarity (lowest *p*-distances) between strain B10 and the strain detected in red-tailed monkey based on the VP6, VP7, NSP3, and NSP4. For the other segments, the B10 strain was genetically more similar to the strains found in primates (including the strain detected in the present study) than to those found in bats. These results further support a simian rather than a bat origin of the B10 strain that was detected in an infant in Kenya.

Finally, in the swab samples of an African palm civet from Inkanamongo, a strain of the genus *Orthoreovirus* (family *Spinareoviridae*) was detected. Orthoreoviruses infect reptiles, birds, and mammals. Currently, ten species of orthoreoviruses are recognised, which are divided into two subgroups based on their fusogenic ability, i.e., their ability to cause fusion of infected cells and, thereby, facilitating a rapid cell-to-cell spread of the infection (Day, 2009): strains of the species *O. mammalis* and *O. piscis* are non-fusogenic, while strains of the other species (i.e., *O. nelsonense, O. papionis*, *O. reptilis*, *O. mahlapitsiense*, *O. broomense*, *O. avis*, and *O. testudinis*) induce the formation of syncytia upon infection.

In the present study, the detected strain phylogenetically clusters with baboon orthoreovirus (genus *Orthoreovirus*, species *O. papionis*) infecting yellow baboons. Baboon orthoreovirus has been detected in the brains of baboons diagnosed with meningoencephalomyelitis (Duncan et al., 1995; Kumar et al., 2014). However, according to the species demarcation criteria proposed for orthoreoviruses (i.e., more than 75% nucleotide sequence identity within a species), the strain detected in the present study represents a new member of the genus *Orthoreovirus* (based only on the RdRp gene sequence), which we tentatively refer to as ‘Civet orthoreovirus’ (Note that strains of orthoreoviruses infecting other carnivores are identified as *O. mammalis* and cluster separate from the strain of ‘Civet orthoreovirus’ (**Supplementary Figure S11**)). Further research is needed to determine the pathogenic and fusogenic potential of this newly detected strain of *Orthoreovirus* and its potential to infect other mammals besides the African palm civet.

The zoonotic potential of a virus is driven by several factors, e.g., a high abundance of the wildlife host, a high prevalence of the virus in the wildlife population, a high frequency of human-wildlife contact, close phylogenetic relationship between the wildlife host and humans, and the ability of a virus to infect a wide taxonomic range of hosts (Wolfe et al., 2007). In this regard, NHPs are of particular concern as a source of zoonotic viral transmission. Non-human primates are most closely related to humans and pose the weakest phylogenetic barrier for interspecies viral transmission. In addition, NHPs are highly prevalent in the African wild meat trade (Kurpiers et al., 2016; Taylor et al., 2015), fostering frequent intimate contact with humans. Indeed, multiple studies report the presence of viruses in NHPs (including the present study) and many viral spillovers from primates to humans have been related to activities involved in the wild meat supply chain, particularly in sub-Saharan Africa (Kurpiers et al., 2016; Morrison-Lanjouw et al., 2023; Smith et al., 2012; Temmam et al., 2017). In the present study, we identified several recognised zoonotic pathogens in NHPs which were genetically closely related to human-infecting strains: a member of the genus *Orthopicobirnavirus* in red-tailed monkey and Allen’s swamp monkey; members of the genus *Simiispumavirus* in red-tailed monkey, Allen’s swamp monkey, and Wolf’s or Dent’s monkey; and a member of the genus *Rotavirus* in red-tailed monkey. The presence of these viruses in the carcasses of these wild mammals highlights the potential public health risk of exposure to these viruses through the intimate human-wildlife interactions related to the wild meat supply chain.

In addition, our study is the first to report on the virome of some mammal species, i.e., tree hyraxes of *Dendrohyrax*, fire-footed and Thomas’s rope squirrel, and Dent’s monkey (although the species identification of the latter is uncertain). Also, we identified known (and unknown) viruses in certain mammal species for the first time (i.e., an orthohepadnavirus in a tree hyrax of the genus *Dendrohyrax*, a kunsagivirus in rope squirrels of the genus *Funisciurus*, orthoherpesviruses in a blue duiker and African palm civet, an orthoreovirus in an African palm civet, a simiispumavirus in Wolf’s or Dent’s monkey, an orthopicobirnavirus in a red-tailed monkey, Allen’s swamp monkey, and northern giant pouched rat, and a rotavirus in a red-tailed monkey), thereby expanding their known host range and genetic diversity. Whether or not these mammals present the natural reservoir or an incidental host of these viruses, as well as the pathogenesis and prevalence, remains an open question. Nevertheless, their ability to harbour these viruses, even for a short period of time, suggests that they may play a role in virus transmission and the emergence of new strains.

It is important to note that the detection of the virus genetic material does not necessarily imply that the virus was still infectious at the time of sampling, especially for cooked and processed meats. However, it is probable that the carcass had still been infectious on several occasions along the supply chain. Furthermore, many viruses can remain viable for a while after the host has been killed (McElhinney et al., 2014; Prescott et al., 2015; Sablone et al., 2021). In sum, our study provides important insights into the diversity of viruses in African wild mammals that were traded for their meat in several regions in the DRC and Belgium, using viral metagenomics. However, the viral diversity that was detected in the present study is probably an underestimation of the true diversity of viruses that infect these hunted mammals. First, the genomic content of the wild meat has been degraded over time depending on their mode of conservation (some samples are older than a decade). For example, samples stored in ethanol at room temperature for over a decade are expected to contain only little intact viral DNA/RNA. Also, degradation of the genomic content by meat processing (e.g., smoking or drying) decreases the chance of viral genome detection (Temmam et al., 2017). Secondly, the detection of viruses is dependent on the sample type as the presence of a virus depends on cellular and tissue tropism, transmission route, and viral properties. Thus, whenever possible, future surveillance efforts should target diverse sample types (i.e., muscle, blood, and various organs and swaps) for each host specimen whenever possible (Mahar et al., 2024). Thirdly, our phylogenetic analyses were limited to viral strains in pools with an RPM greater than 1, which likely underestimate the number of virus-positive pools. This is evidenced by the detection of a few SIV reads in pool 51, which tested positive for SIV using a targeted PCR assay. This underscores the high sensitivity of targeted PCR assays once the presence of a specific virus is established and highlights the necessity for additional PCR testing on pools with an RPM below 1. Moreover, to determine the infected individual within a pool and identify the potential presence of multiple viral strains, further PCR assays and Sanger sequencing should be conducted on unpooled samples. Primers for these assays can be designed based on the viral consensus sequences characterized in the present study. Finally, our metagenomic analyses did not always result in the recovery of the entire viral genome. Deeper sequencing or additional approaches, such as PCR using strain-specific primers to fill gaps in the genome, could be used to determine the full viral genome, as illustrated by e.g. Bletsa et al. (2021).

## DECLARATION OF COMPETING INTEREST

The authors declare that they have no competing interests.

## DATA AVAILABILITY

The sequences generated in this study will be submitted to the GenBank repository upon acceptance.

## AUTHOR CONTRIBUTIONS

SGr and PG supervised and coordinated the study. SGr, SGo, EV, PG, and AC contributed to the conception and design of the study. CN, DoA, DuA, PB, GCG, LJ, AL, NL, HL, CM, JM, SN, RV, and EV participated in the sample collection. The laboratory work and mammal species identification was conducted by JT, SGo, AV and SGr. MG conducted the metagenomic analyses and wrote the first draft of the manuscript. All authors revised the draft and approved the final manuscript.

## FUNDING

This work was funded by the World Wildlife Fund (France) (SGr), ERA-NET BiodivERsA— EC funding BIODIV-AFREID (Research Foundation—Flanders: G0G2119N) (HL, EV, DuA), Flemish Interuniversity Council South-Initiative (VLIR-UOS SI) (AL, DuA, EV), the Flemish Interuniversity Council (VLIR-CUI) (DuA, EV), the Research Foundation—Flanders (Fundamental Research Project G051322N), and a FED-tWIN scholar funded by the Belgian federal government (Prf-2019-prf004_OMEga) (SGr). We thank the Capacities for Biodiversity and Sustainable Development (CEBioS) program, based at the Royal Belgian Institute of Natural Sciences and funded by the Belgian Development Cooperation (DGD), for their financial support to the CSB team members. The Barcoding Facility for Tissues and Organisms of Policy Concern (BopCo) was funded by Belspo (RT23BopCo-CE).

## ACKNOWLEDGEMENTS

We appreciate the help of all people involved in the field work and sampling procedure. Furthermore, we would like to thank Natalie Van Houtte for her technical support in the laboratory.

## ETHICAL CLEARANCE AND PERMITS

For compliance with ethical standards for the samples collected at African grocery stores in Brussels, Belgium, we refer to Gombeer et al. (2021). For permits and ethical clearance of samples collected in the Tshuapa province, DRC, in 2021, we refer to van Vredendaal et al. (2024). Samples from the archived collection of the RBINS and UAntwerp were collected between 2010 and 2014, before the Nagoya protocol was formally implemented in the DRC. However, material transfer agreements (MTA) for these samples were provided by the University of Kisangani’s Biodiversity Surveillance Center (CSB). We did not apply for a CITES export permit as the identity of the meat (i.e., mammal species) was unknown at the time.

## Notes

### Competing Interest Statement

The authors have declared no competing interest.

