## Supplementary figures for "A wide diversity of viruses detected in African mammals involved in the wild meat supply chain"

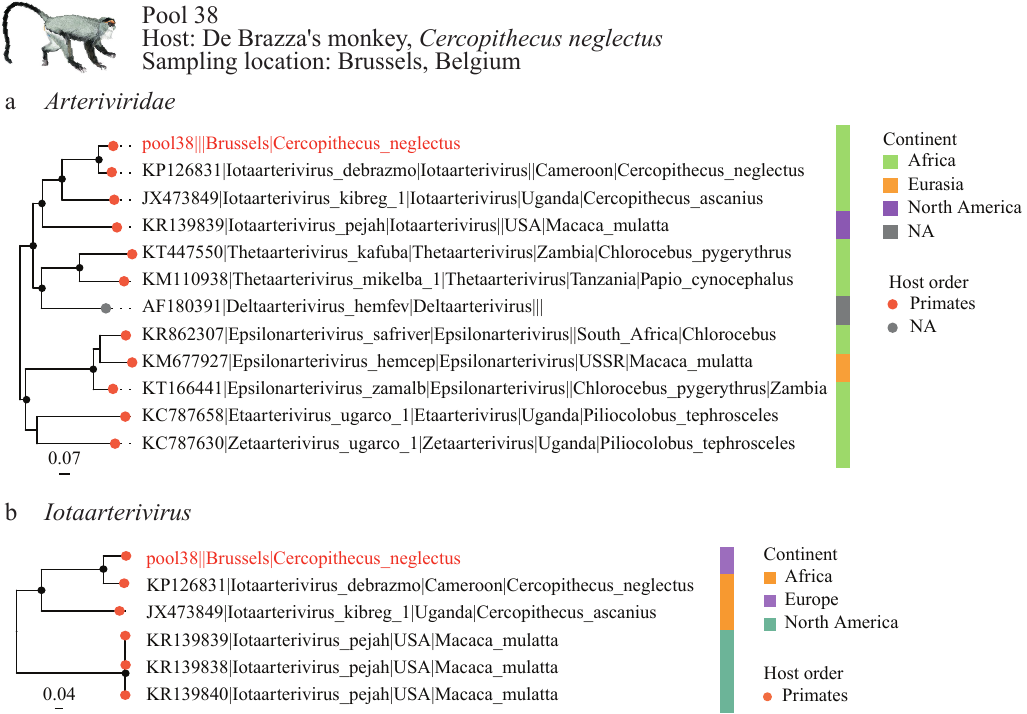

**Supplementary Figure S1. Phylogenetic relationships of members of *Arteriviridae* (a) and *Iotaarterivirus* (b) inferred from the RNA-dependent RNA polymerase gene.** Maximum likelihood phylogenetic trees with only well-supported nodes (bootstrap ≥ 85 %) indicated by black dots. Scale bar indicates the mean number of nucleotide substitutions per site. For each strain that is included in the analysis, the GenBank accession (or pool) number, virus name, genus (for (a)), country, and host are given (when available from NCBI Virus). Tip points are coloured by the host order (humans indicated by a different colour than other primates), while squares next to each tip label are coloured according to the continent. The viral strain detected in the present study is in red. Note that the continent represents the location of sampling and does not imply that samples were collected from wild animals. The origin of the samples that were collected in Brussels in the present study were assigned to Africa as the meat was imported from this continent. Mammal image of the host is from Kingdon (2015).

**Supplementary Figure S2**

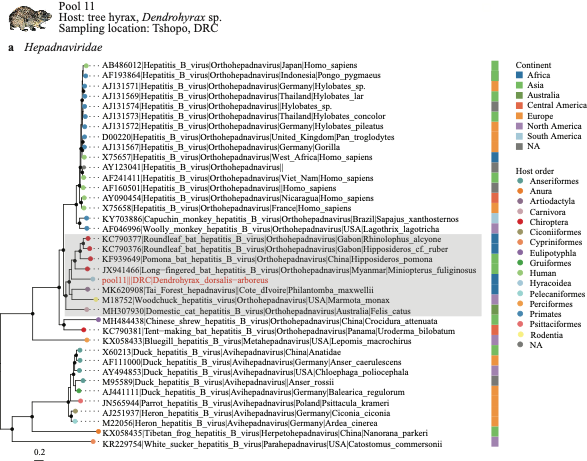

**Supplementary Figure S2. Continued**

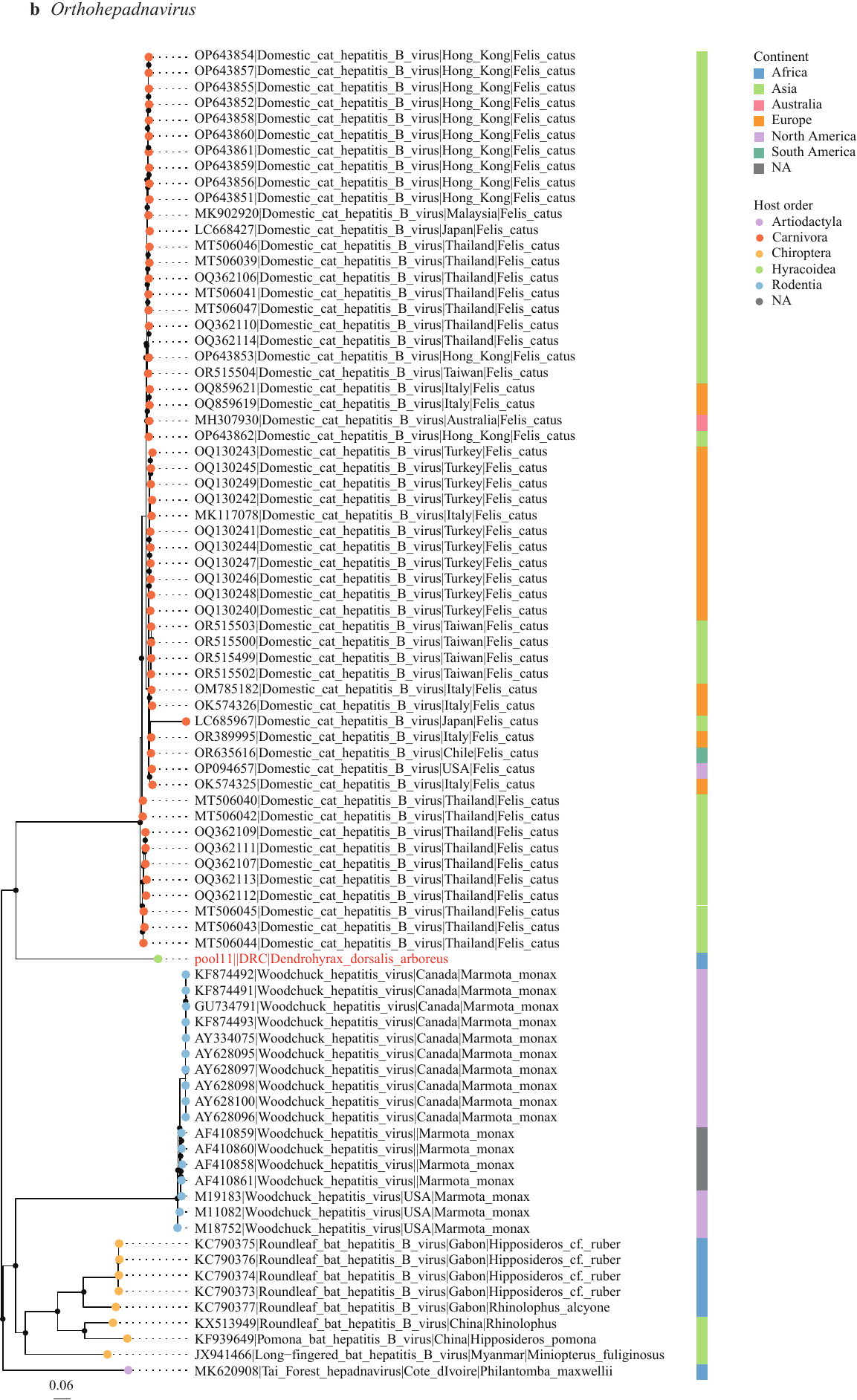

**Supplementary Figure S2. Phylogenetic relationships of members of *Hepadnaviridae* (a) and *Orthohepadnavirus* (b) inferred from the DNA polymerase gene.** Maximum likelihood phylogenetic trees with only well-supported nodes (bootstrap ≥ 85 %) indicated by black dots. Scale bar indicates the mean number of nucleotide substitutions per site. For the phylogenetic tree of *Orthohepadnavirus* (b) we only included specimens of species that phylogenetically clustered in the same clade as the detected virus in the family-level phylogeny (framed in grey in (a)) to maintain a manageable dataset. For each strain that is included in the analysis, the GenBank accession (or pool) number, virus name, genus (for (a)), country, and host are given (when available from NCBI Virus). Tip points are coloured by the host order (humans indicated by a different colour than other primates), while squares next to each tip label are coloured according to the continent. The viral strain detected in the present study is in red. Note that the continent represents the location of sampling and does not imply that samples were collected from wild animals. Mammal image of the host (we chose *Dendrohyrax arboreus* to depict *D.* sp.) is from Kingdon (2015).

**Supplementary Figure S3**

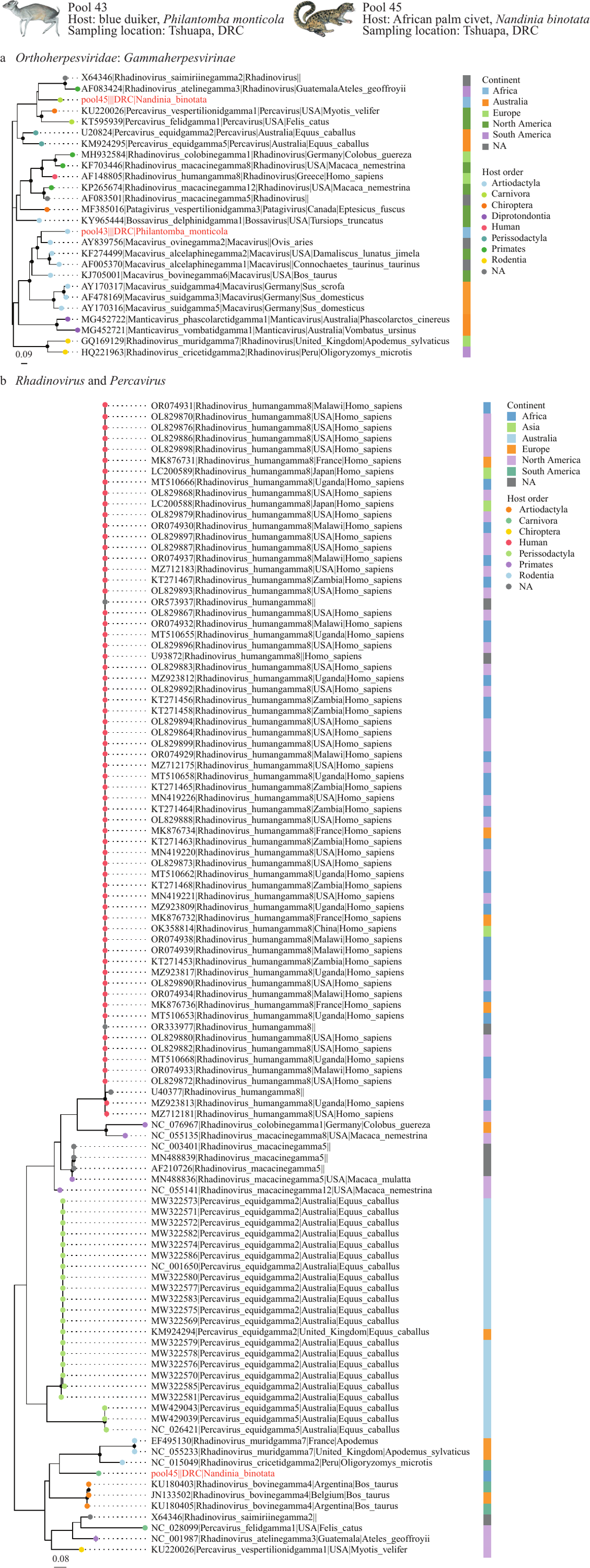

**Supplementary Figure S3. Continued**

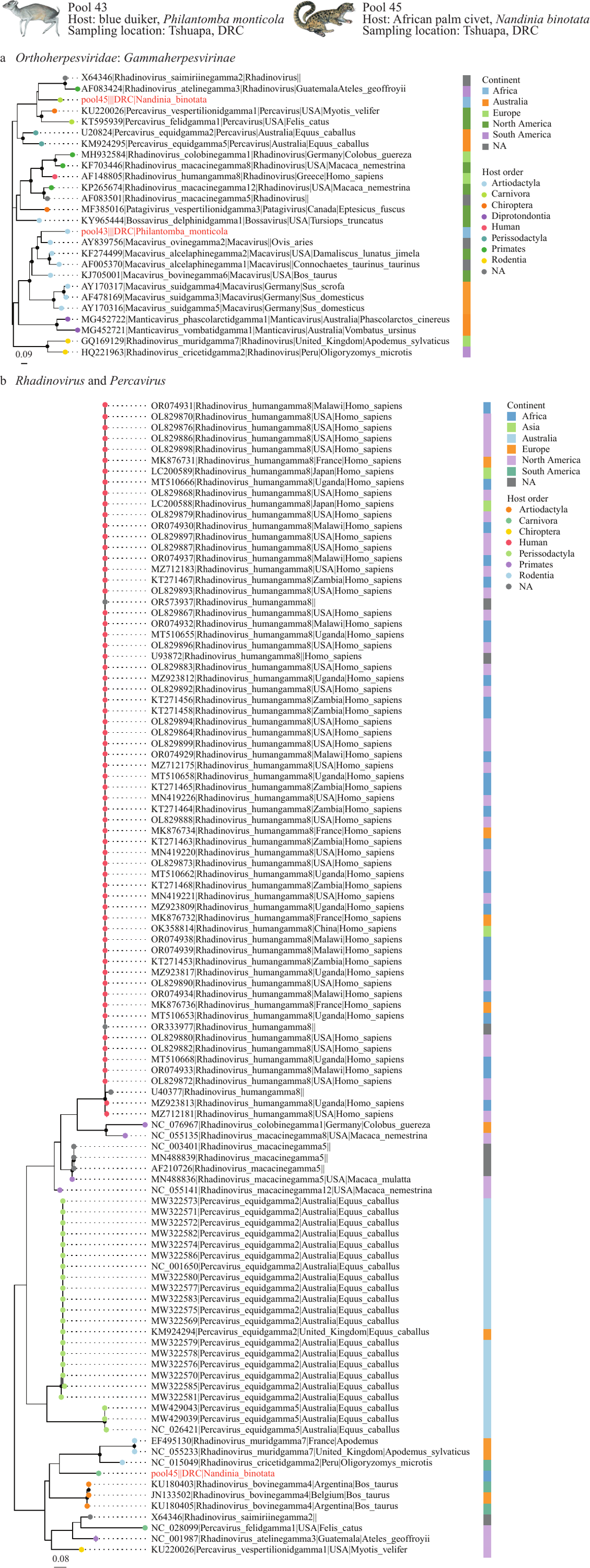

**Supplementary Figure S3.** **Phylogenetic relationships of members of *Gammaherpesvirinae* (*Orthoherpesviridae*) (a) and *Rhadinovirus* and *Percavirus* (b) inferred from the capsid gene.** Maximum likelihood phylogenetic trees with only well-supported nodes (bootstrap ≥ 85 %) indicated by black dots. Scale bar indicates the mean number of nucleotide substitutions per site. For each strain that is included in the analysis, the GenBank accession (or pool) number, virus name, genus (for (a)), country, and host are given (when available from NCBI Virus). Tip points are coloured by the host order (humans indicated by a different colour than other primates), while squares next to each tip label are coloured according to continent. The viral strain detected in the present study in red. Note that the continent represents the location of sampling and does not imply that samples were collected from wild animals. Mammal images of the hosts are from Kingdon (2015).

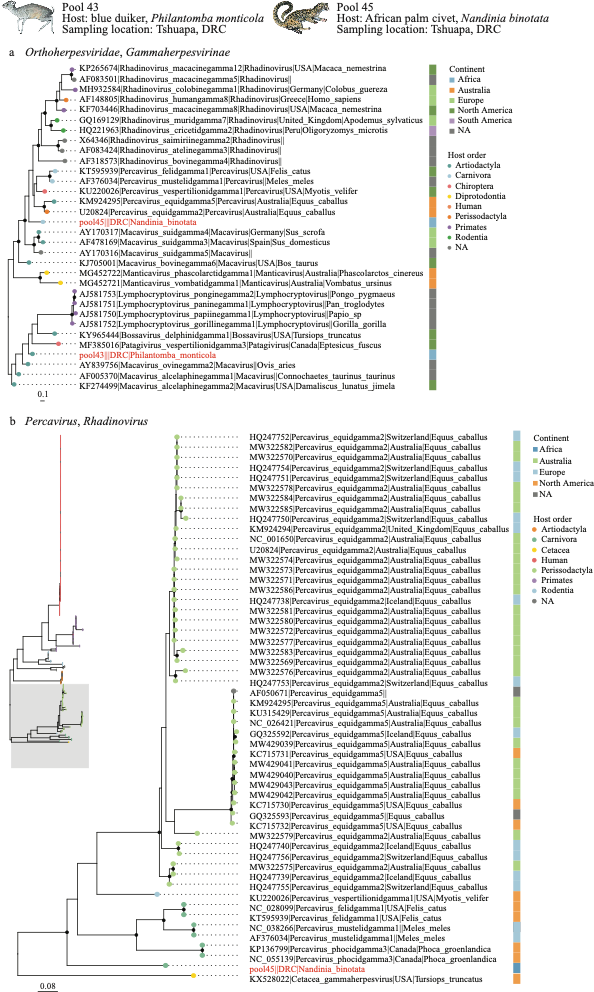

**Supplementary Figure S4. Phylogenetic relationships of members of *Gammaherpesvirinae* (*Orthoherpesviridae*) (a) and *Rhadinovirus* and *Percavirus* (b) inferred from the envelope glycoprotein B gene.** Maximum likelihood phylogenetic trees with only well-supported nodes (bootstrap ≥ 85 %) indicated by black dots. Scale bar indicates the mean number of nucleotide substitutions per site. For each strain that is included in the analysis, the GenBank accession (or pool) number, virus name, genus (for (a)), country, and host are given (when available from NCBI Virus). Tip points are coloured by the host order (humans indicated by a different colour than other primates), while squares next to each tip label are coloured according to the continent. The viral strain detected in the present study is in red. In (b), a subset of the entire phylogenetic tree (framed in grey at the left) is shown magnified (tip labels and support values are omitted in the entire phylogenetic tree to increase readability). Note that the continent represents the location of sampling and does not imply that samples were collected from wild animals. Mammal images of the hosts are from Kingdon (2015).

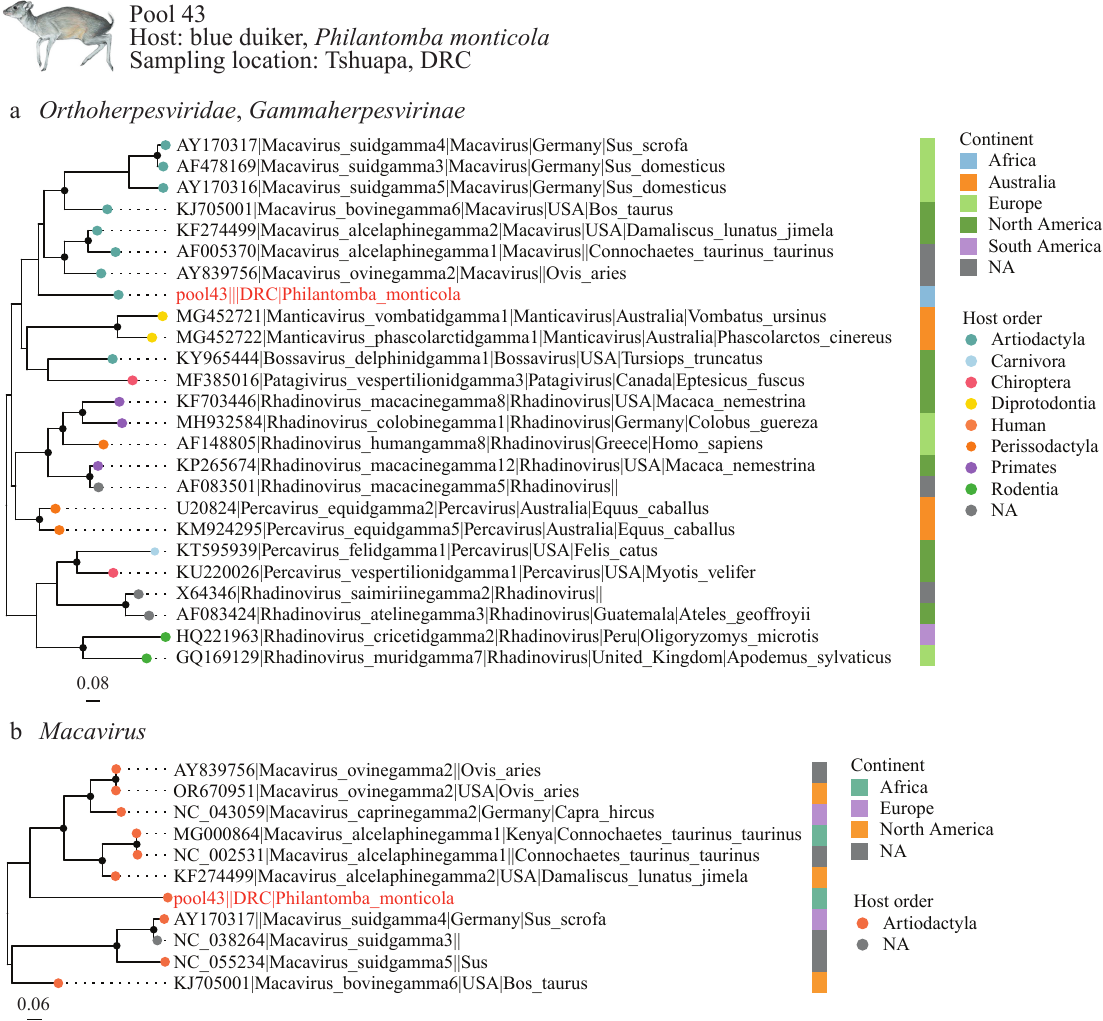

**Supplementary Figure S5. Phylogenetic relationships of members of *Gammaherpesvirinae* (*Orthoherpesviridae*) (a) and *Macavirus* (b) inferred from the DNA polymerase gene.** Maximum likelihood phylogenetic trees with only well-supported nodes (bootstrap ≥ 85 %) indicated by black dots. Scale bar indicates the mean number of nucleotide substitutions per site. For each strain that is included in the analysis, the GenBank accession (or pool) number, virus name, genus (for (a)), country, and host are given (when available from NCBI Virus). Tip points are coloured by the host order (humans indicated by a different colour than other primates), while squares next to each tip label are coloured according to the continent. The viral strain detected in the present study is in red. Mammal images of the host is from Kingdon (2015).

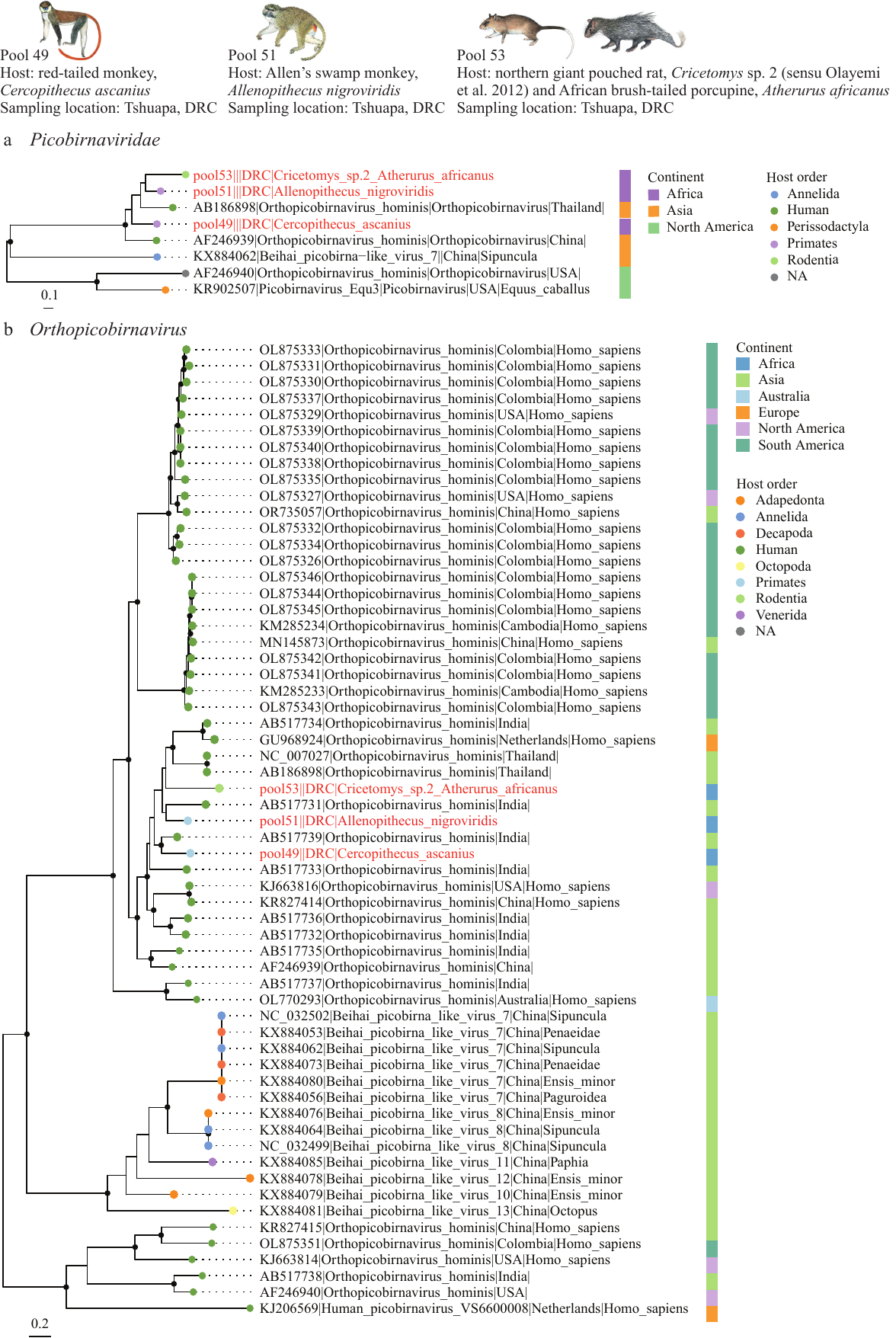

**Supplementary Figure S6. Phylogenetic relationships of members of *Picobirnaviridae* (a) and *Orthopicobirnavirus* (b) inferred from the RNA-dependent RNA polymerase gene.** Maximum likelihood phylogenetic trees with only well-supported nodes (bootstrap ≥ 85 %) indicated by black dots. Scale bar indicates the mean number of nucleotide substitutions per site. For each strain that is included in the analysis, the GenBank accession (or pool) number, virus name, genus (for (a)), country, and host are given (when available from NCBI Virus). Tip points are coloured by the host order (humans indicated by a different colour than other primates), while squares next to each tip label are coloured according to the continent. The viral strain detected in the present study is in red. Note that the continent represents the location of sampling and does not imply that samples were collected from wild animals. Mammal images of the hosts are from Kingdon (2015). We chose *Cricetomys gambianus* as a depiction of *C.* sp. 2 (sensu Olayemi et al., 2012).

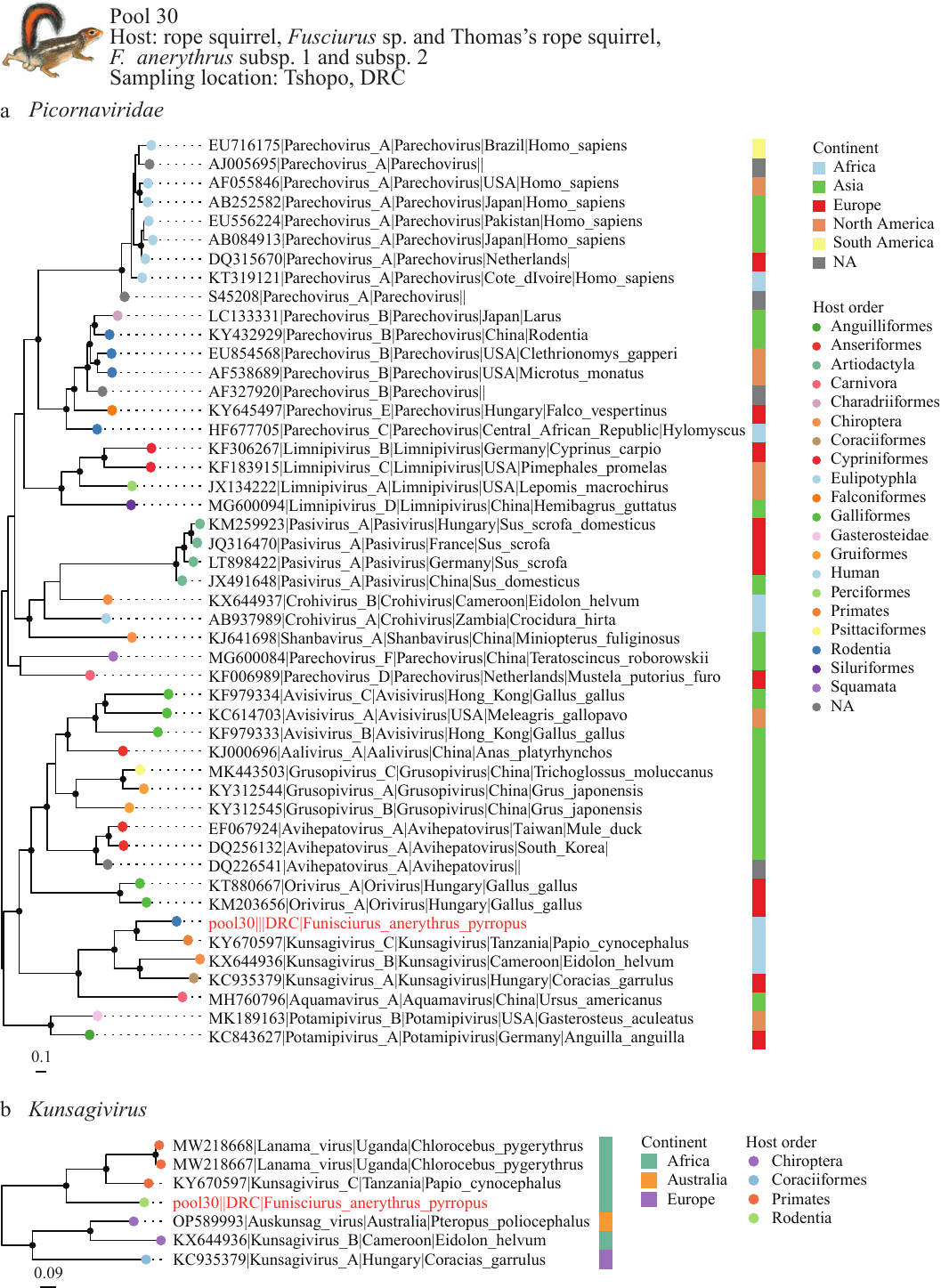

**Supplementary Figure S7. Phylogenetic relationships of members of *Picornaviridae* (a) and *Kunsagivirus* (b) inferred from the RNA-dependent RNA polymerase gene.** Maximum likelihood phylogenetic trees with only well-supported nodes (bootstrap ≥ 85 %) indicated by black dots. Scale bar indicates the mean number of nucleotide substitutions per site. For each strain that is included in the analysis, the GenBank accession (or pool) number, virus name, genus (for (a)), country, and host are given (when available from NCBI Virus). Tip points are coloured by the host order (humans indicated by a different colour than other primates), while squares next to each tip label are coloured according to the continent. The viral strain detected in the present study is in red. Note that the continent represents the location of sampling and does not imply that samples were collected from wild animals. Mammal image of the host (*F. anerythrus*) is from Kingdon (2015).

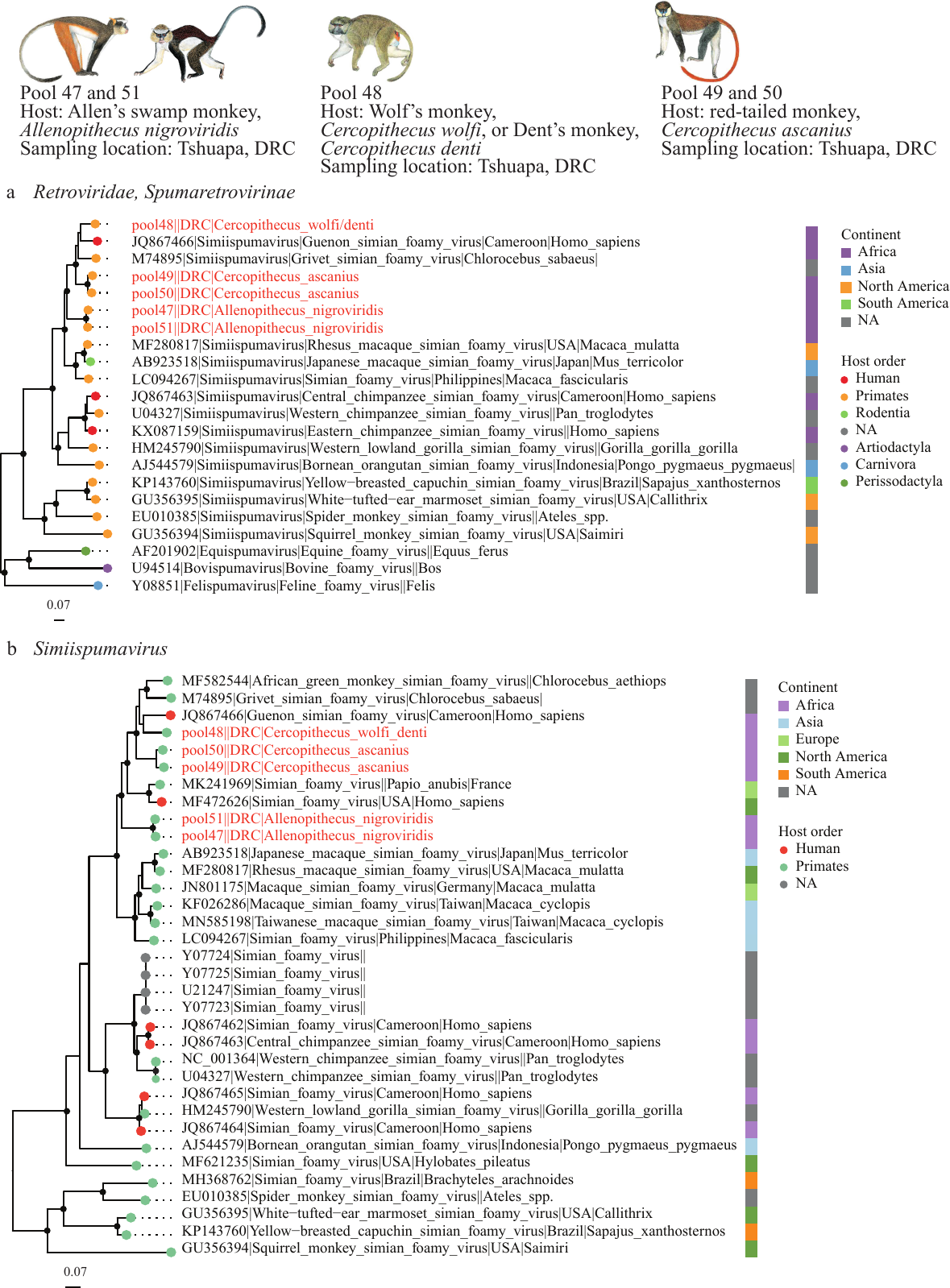

**Supplementary Figure S8. Phylogenetic relationships of members of *Retroviridae* (a) and *Simiispumavirus* (b) inferred from the RNA-dependent RNA polymerase gene.** Maximum likelihood phylogenetic trees with only well-supported nodes (bootstrap ≥ 85 %) indicated by black dots. Scale bar indicates the mean number of nucleotide substitutions per site. For each strain that is included in the analysis, the GenBank accession (or pool) number, virus name, genus (for (a)), country, and host are given (when available from NCBI Virus). Tip points are coloured by the host order (humans indicated by a different colour than other primates), while squares next to each tip label are coloured according to the continent. The viral strain detected in the present study is in red. Note that the continent represents the location of sampling and does not imply that samples were collected from wild animals. Mammal images of the hosts are from Kingdon (2015).

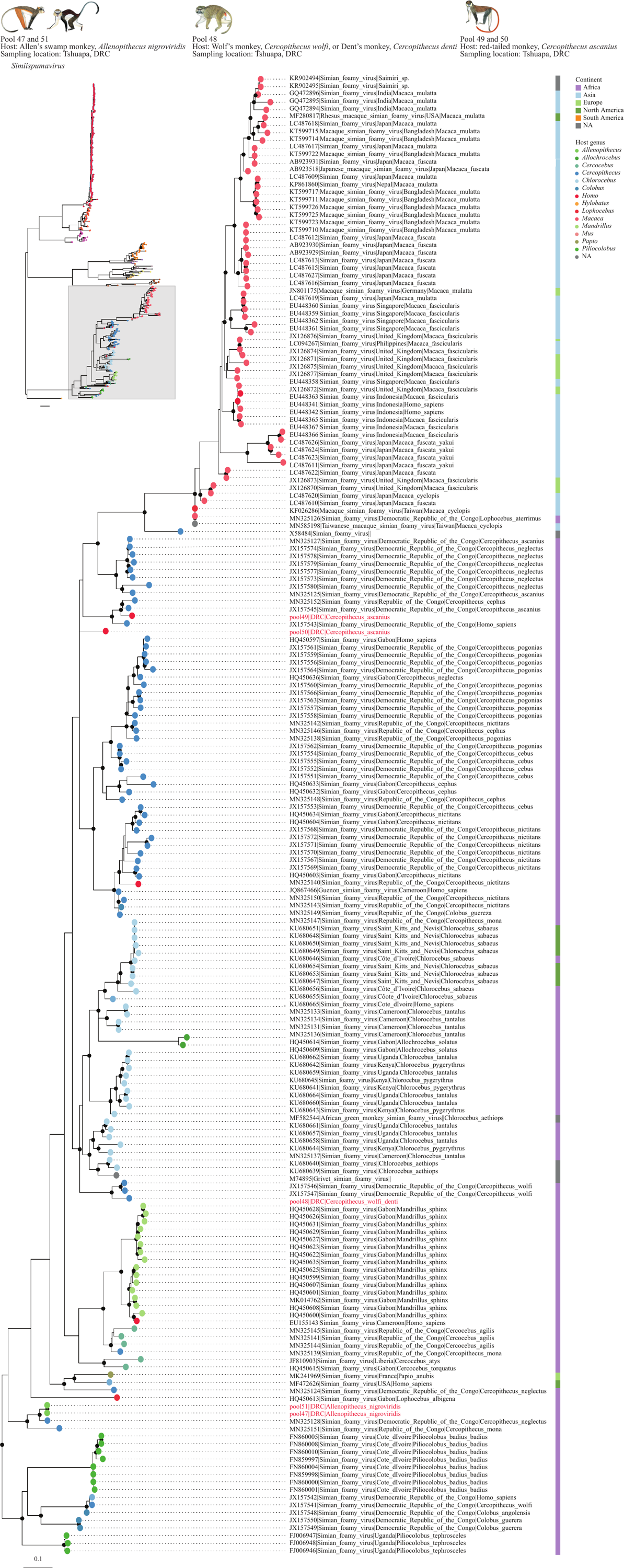

**Supplementary Figure S9. Phylogenetic relationships of members of *Simiispumavirus* inferred from a short (271 bp) fragment of the RNA-dependent RNA polymerase gene.** Maximum likelihood phylogenetic trees with only well-supported nodes (bootstrap ≥ 85 %) indicated by black dots. Scale bar indicates the mean number of nucleotide substitutions per site. For each strain that is included in the analysis, the GenBank accession (or pool) number, virus name, country, and host are given (when available from NCBI Virus). Tip points are coloured by the primate. genus, while squares next to each tip label are coloured according to the continent. The viral strain detected in the present study is in red. Note that the continent represents the location of sampling and does not imply that samples were collected from wild animals. Mammal images of the hosts are from Kingdon (2015).

**Supplementary Figure S10**

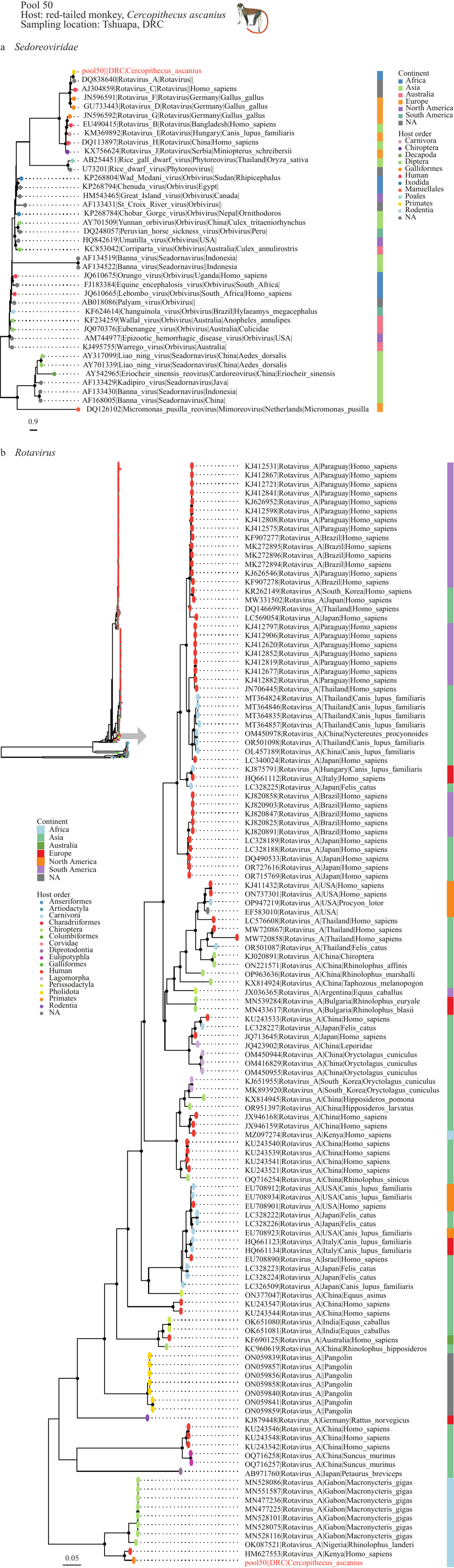

**Supplementary Figure S10. Continued**

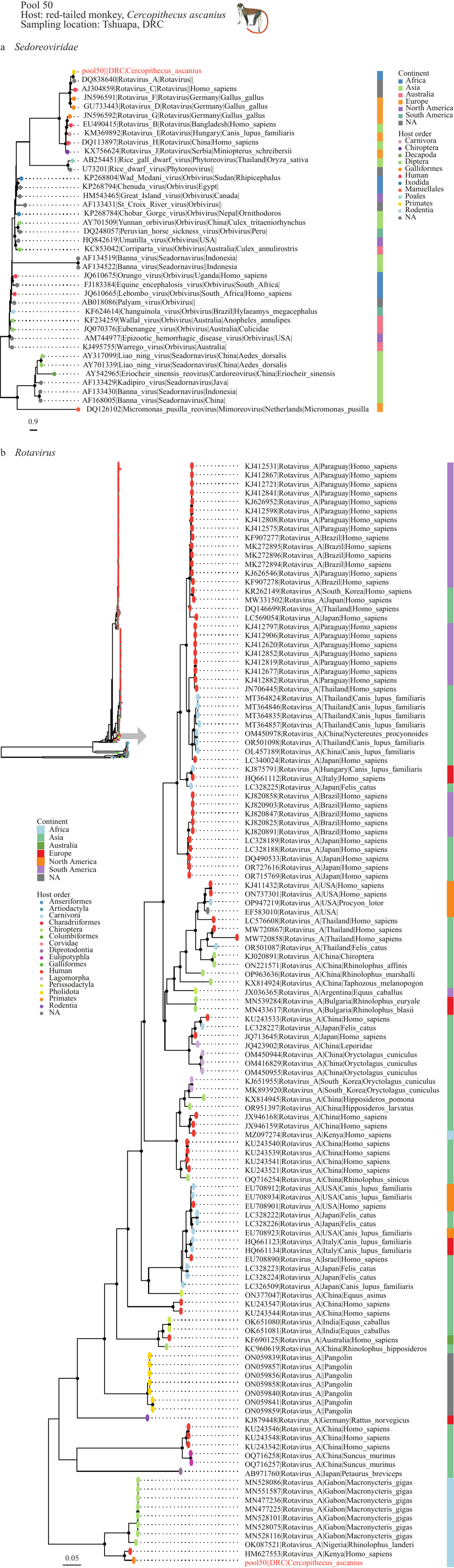

**Supplementary Figure S10. Phylogenetic relationships of members of *Sedoreoviridae* (a) and *Rotavirus* (b) inferred from RNA-dependent RNA polymerase.** Maximum likelihood phylogenetic trees with only well-supported nodes (bootstrap ≥ 85 %) indicated by black dots. Scale bar indicates the mean number of nucleotide substitutions per site. For each strain that is included in the analysis, the GenBank accession (or pool) number, virus name, genus (for (a)), country, and host are given (when available from NCBI Virus). Tip points are coloured by the host order (humans indicated by a different colour than other primates), while squares next to each tip label are coloured according to the continent. The viral strain detected in the present study is in red. In (b) a subset of the entire phylogenetic tree (marked with a grey arrow at the left) is shown magnified (support values and tip labels are omitted in the entire phylogenetic tree to increase readability). Note that the continent represents the location of sampling and does not imply that samples were collected from wild animals. Mammal images of the hosts are from Kingdon (2015).

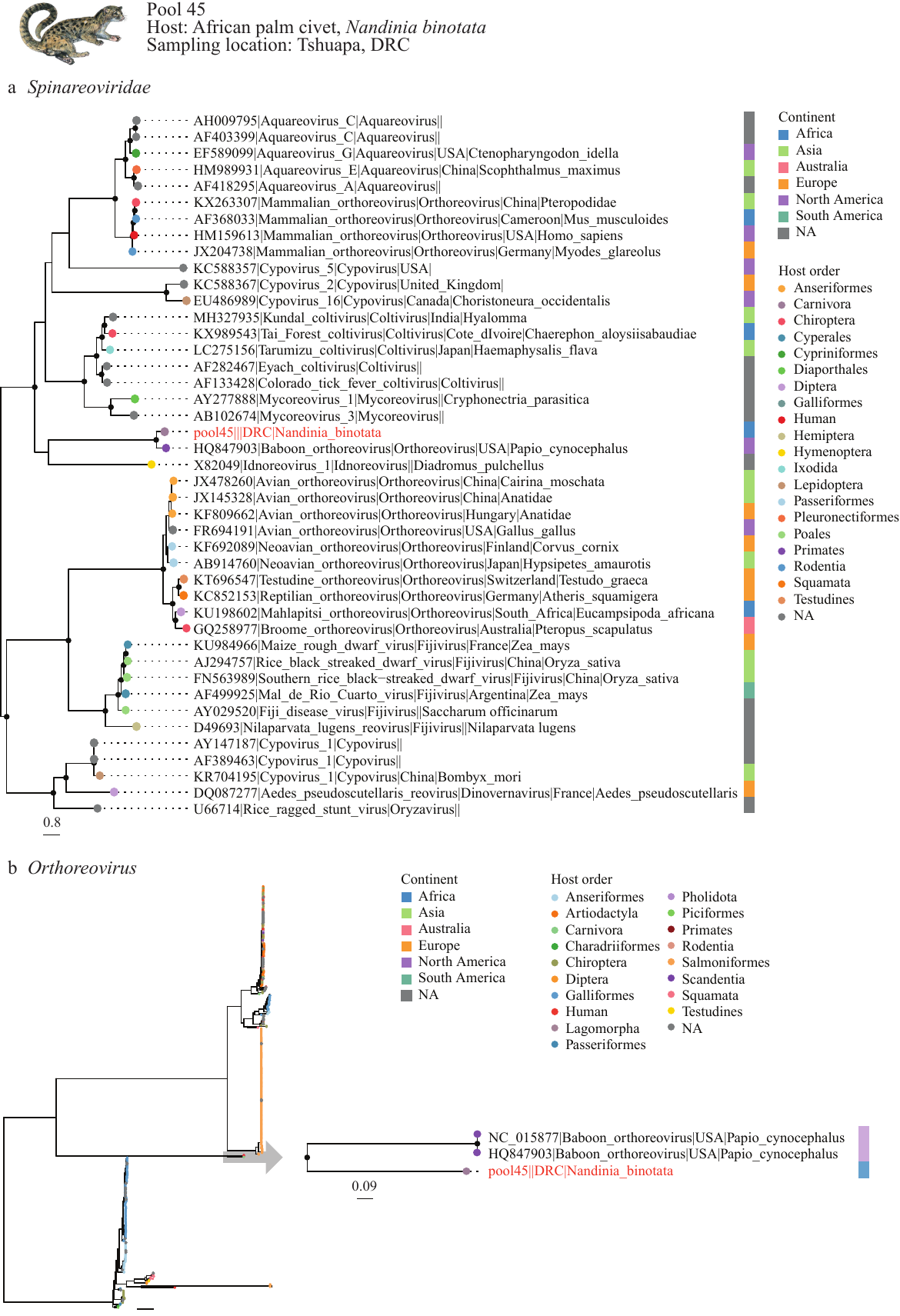

**Supplementary Figure S11. Phylogenetic relationships of members of *Spinareoviridae* (a) and *Orthoreovirus* (b) inferred from RNA-dependent RNA polymerase.** Maximum likelihood phylogenetic trees with only well-supported nodes (bootstrap ≥ 85 %) indicated by black dots. Scale bar indicates the mean number of nucleotide substitutions per site. For each strain that is included in the analysis, the GenBank accession (or pool) number, virus name, genus (for (a)), country, and host are given (when available from NCBI Virus). Tip points are coloured by the host order (humans indicated by a different colour than other primates), while squares next to each tip label are coloured according to the continent. The viral strain detected in the present study is in red. In (b) a subset of the entire phylogenetic tree (marked with a grey arrow at the right) is shown magnified (support values and tip labels are omitted in the entire phylogenetic tree to increase readability). Note that the continent represents the location of sampling and does not imply that samples were collected from wild animals. Mammal images of the hosts are from Kingdon (2015).
